# Synaptic accumulation of FUS triggers age-dependent misregulation of inhibitory synapses in ALS-FUS mice

**DOI:** 10.1101/2020.06.10.136010

**Authors:** Sonu Sahadevan, Katharina M. Hembach, Elena Tantardini, Manuela Pérez-Berlanga, Marian Hruska-Plochan, Julien Weber, Petra Schwarz, Luc Dupuis, Mark D. Robinson, Pierre De Rossi, Magdalini Polymenidou

## Abstract

FUS is a primarily nuclear RNA-binding protein with important roles in RNA processing and transport. FUS mutations disrupting its nuclear localization characterize a subset of amyotrophic lateral sclerosis (ALS-FUS) patients, through an unidentified pathological mechanism. FUS regulates nuclear RNAs, but its role at the synapse is poorly understood. Here, we used super-resolution imaging to determine the physiological localization of extranuclear, neuronal FUS and found it predominantly near the vesicle reserve pool of presynaptic sites. Using CLIP-seq on synaptoneurosome preparations, we identified synaptic RNA targets of FUS that are associated with synapse organization and plasticity. Synaptic FUS was significantly increased in a knock-in mouse model of ALS-FUS, at presymptomatic stages, accompanied by alterations in density and size of GABAergic synapses. RNA-seq of synaptoneurosomes highlighted age-dependent dysregulation of glutamatergic and GABAergic synapses. Our study indicates that FUS accumulation at the synapse in early stages of ALS-FUS results in synaptic impairment, potentially representing an initial trigger of neurodegeneration.

## Introduction

FUS (Fused in sarcoma) is a nucleic acid binding protein involved in several processes of RNA metabolism^1^. Physiologically, FUS is predominantly localized to the nucleus^2^ via active transport by transportin (TNPO)^3^ and it can shuttle to the cytoplasm by passive diffusion^4,5^. In amyotrophic lateral sclerosis (ALS) and frontotemporal dementia (FTD), FUS mislocalizes to the cytoplasm where it forms insoluble aggregates^6–8^. In ALS, cytoplasmic mislocalization of FUS is associated with mutations that are mainly clustered in the proline-tyrosine nuclear localization signal (PY-NLS) at the C-terminal site of the protein^9^ and lead to mislocalization of the protein to the cytosol. However, in FTD, FUS mislocalization occurs in the absence of mutations^10^. FUS is incorporated in cytoplasmic stress granules^5,11^ and undergoes concentration-dependent, liquid-liquid phase separation^12,13^, which is modulated by binding of TNPO and arginine methylation of FUS^14–17^. This likely contributes to the role of FUS in forming specific identities of ribonucleoprotein (RNP) granules^18,19^ and in transporting RNA cargos^20^, which is essential for local translation in neurons^21^.

Despite the central role of FUS in neurodegenerative diseases, little is known about its function in specialized neuronal compartments, such as synapses. FUS was shown to mediate RNA transport^20^ and is involved in stabilization of RNAs that encode proteins with important synaptic functions^22^, such as *GluA1* and *SynGAP1*^23,24^. While the presence of FUS protein in synaptic compartments has been confirmed, its exact subsynaptic localization is debated. Diverging results described the presence of FUS at the pre-synapses in close proximity to synaptic vesicles^25–27^, but also in dendritic spines^20^ and in association with the postsynaptic density^28^. Confirming a functional role of FUS at the synaptic sites, behavioral and synaptic morphological changes have been observed upon depletion of FUS in mouse models^23,29,30^. Notably, mouse models associated with mislocalization of FUS exhibited reduced axonal translation contributing to synaptic impairments^31^. Synaptic dysfunction has been suggested to be the early event of several neurodegenerative disorders including ALS and FTD^32–36^. The disruption of RNA-binding proteins (RBPs) and RNA regulation could be a central cause of synaptic defects in these disorders.

Previous studies identified nuclear RNA targets of FUS with different cross-linking immunoprecipitation and high-throughput sequencing (CLIP-seq) approaches^22,37–41^. Collectively, these findings showed that FUS binds mainly introns, without a strong sequence specificity, but a preference for either GU-rich regions^22,38,40,41^, which is mediated via its zinc finger (ZnF) domain, or a stem-loop RNA^37^ via its RNA recognition motif^42^. FUS often binds close to alternatively spliced exons, highlighting its role in splicing regulation^22,38,39^. CLIP-seq studies also identified RNAs bound by FUS at their 3’ untranslated regions (3’UTRs) and exons^22,39,41^, suggesting a direct role of FUS in RNA transport and regulating synaptic mRNA stability^23,24^ and polyadenylation^40^. However, a precise list of synaptic RNAs directly regulated by FUS is yet to be identified.

In this study, we focused on understanding the role of synaptic FUS in RNA homeostasis and the consequences of ALS-causing mutations in FUS on synaptic maintenance. Using super-resolution imaging, we confirmed the presence of FUS at the synapse. FUS was found at both excitatory and inhibitory synapses, was enriched at the presynapse and rarely associated with postsynaptic structures. Synaptoneurosome preparations from adult mouse cortex, coupled with CLIP-seq uncovered specific synaptic RNA targets of FUS. Computational analyses revealed that most of these targets were associated with both glutamatergic and GABAergic networks. In a heterozygous knock-in FUS mouse model, which harbors a deletion in the NLS of FUS allele, thereby mimicking the majority of ALS-causing mutations^43^, we found significant increase of synaptic FUS localization. To test the effect of this elevation in synaptic FUS, we investigated the synaptic organization of the hippocampus, which is enriched in glutamatergic and GABAergic synapses, and found mild and transient changes. However, RNA-seq analysis revealed age-dependent alterations of synaptic RNA composition including glutamatergic and GABAergic synapses. Our data indicate that early synaptic alterations in the GABAergic network precede motor impairments in these ALS-FUS mice^43^, and may trigger early behavioral dysfunctions, such as hyperactivity and social disinhibition that these mice develop (Scekic-Zahirovic, Sanjuan-Ruiz et al., co-submitted manuscript).

Altogether, our results demonstrate a critical role for FUS in synaptic RNA homeostasis via direct association with specific synaptic RNAs, such as *Gabra1, Grin1* and others. Our study indicates that enhanced synaptic localization of FUS in early stages of ALS-FUS results in synaptic impairment, potentially representing the initial trigger of neurodegeneration. Importantly, we show that increased localization of FUS at the synapses, in the absence of aggregation, suffices to cause synaptic impairment.

## Results

### FUS is enriched at the presynaptic compartment of mature cortical and hippocampal neurons

While FUS has been shown at synaptic sites, its exact subsynaptic localization is debated. Some studies described a presynaptic enrichment of FUS in cortical neurons and motoneurons^25,27^, whereas others have shown an association of FUS with postsynaptic density (PSD) sites^20,28^. To clarify the precise localization of FUS at the synapses, we first performed confocal analysis in mouse cortex (**Fig. 1a-b**) and hippocampus (**Supplementary Fig. 1a-b**), which confirmed the presence of extranuclear FUS clusters along dendrites and axons (identified with MAP2 and PNF, respectively) and associated with synaptic markers (Synapsin1 and PSD95). To determine the precise subsynaptic localization of FUS, we used super-resolution microscopy (SRM) imaging of mouse hippocampal and cortical synapses. We first explored the distribution of FUS between excitatory and inhibitory synapses of cortical and hippocampal neurons (**Fig. 1c**). STED (Stimulated emission depletion) microscopy was used to precisely determine the localization of FUS clusters compared to synaptic markers: VGAT was used as a marker for inhibitory synapses and PSD95 for excitatory synapses. Image analysis was used to calculate the distance of the closest neighbor (**Supplementary Fig. 1c**). Only FUS clusters within 200 nm from a synaptic marker were considered for this analysis. Our results showed that extranuclear FUS preferentially associates with excitatory synapses, with 46% of the detected ones containing FUS, while only 20% of analyzed inhibitory synapses showing FUS positivity (t-test, p=0.0016) (**Fig. 1d**).

**Fig. 1.**
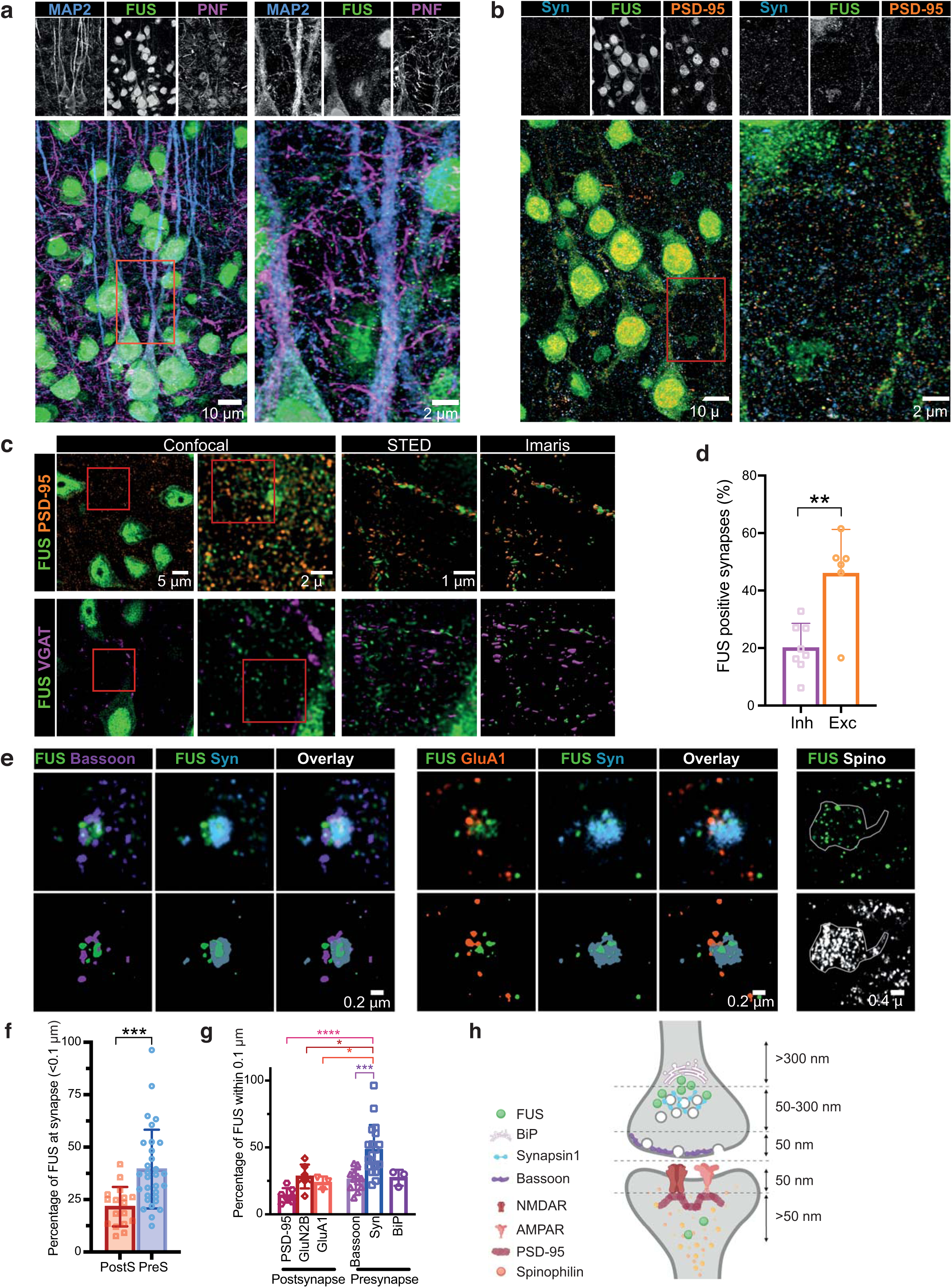
FUS is enriched at the presynaptic compartment. **(a)** Confocal images showing the distribution of FUS (green) in the pyramidal layer of the retrosplenial cortical area along with MAP2 (blue) and PNF (magenta). Left panel shows the overview and the right panel the zoomed in area labelled with the red box on the left panel. **(b)** Similar confocal images showing FUS (green) along with PSD95 (orange) and Synapsin 1 (Syn, blue. (**c**) Synaptic localization of FUS was assessed by STED microscopy using excitatory (PSD95) and inhibitory (VGAT) markers for synapses. 60 μm brain sections were analyzed and distance between FUS and the synaptic markers was analyzed using Imaris. (**d**) Bar graph representing the percentage of synapses within 200 nm of FUS clusters and showing an enrichment of FUS at the excitatory synapses. (**e**) dSTORM was used to explore more precisely the FUS localization within the synapse, using primary culture. Bassoon and Synapsin 1 (Syn) were used to label the presynaptic compartment and GluN1, GluA1 and PSD95 were used to label the postsynapse. Spinophilin (Spino) was used to label the spines. (**f**) Bar graph representing the percentage of FUS localized within 100nm from presynaptic or postsynaptic markers. (**g**) Bar graph representing the distribution of FUS in the synapse. (**h**) Schematic summarizing the FUS localization within the synapse. Graph bar showing mean + SD. *p>0.05, **p>0.01, ***p>0.001, ****p>0.000.

To better define the precise localization of FUS within the synapse, cortical and hippocampal primary cultures were immunolabeled for FUS along with pre- and postsynaptic markers (**Fig. 1e** and **Supplementary Fig. 1d-e**) and their relative distance was analyzed. At the presynapse, Synapsin 1 was used to label the vesicle reserve pool^44^, and Bassoon to label the presynaptic active zone^45^. At the postsynaptic site, GluN2B, subunit of NMDA receptors, and GluA1, subunit of AMPA receptors, were used to label glutamatergic synapses. PSD95 was used to label the postsynaptic density zone^46^. Distribution of FUS at the synapse showed a closer association with Synapsin 1 compared to Bassoon, GluA1, BiP (ER marker) and GluN2B (**Supplementary Fig. 1f-g**). FUS also appeared to be closer to Bassoon compared to PSD95 (**Supplementary Fig. 1f-g**). A subset of FUS was also localized at the spine (**Fig. 1e**). To strengthen our analyses and to refine the precise localization of FUS, the relative proportion of FUS within 100 nm was compared for each marker. Our results showed a preferential FUS localization at the presynaptic site (**Fig. 1f**) (t-test, p=0.0006), in accordance with previously reported data^25,27^. Within the presynaptic site (**Fig. 1g**), FUS was significantly enriched in the Synapsin-positive area (One-way ANOVA, p<0.0001, posthoc Tukey, Syn1 vs. PSD95, p<0.0001; Syn1 vs. GluN2B, p=0.0157; Syn1 vs. GluA1, p=0.454; Syn1 vs. Bassoon, p=0.0005). However, no significant difference was found with the ER marker, suggesting that FUS could be localized between Synapsin 1 and ER at the presynapse (**Fig. 1h**). These results are in line with the previously published localization of FUS within 150 nm from the active presynaptic zone^27^, but highlight the presence of FUS also at the postsynaptic site, potentially explaining the apparently contradictory results of previous studies^20,28^.

### Identification of synaptic RNA targets of FUS

The role of FUS in the nucleus has been well studied and previously published CLIP-seq data identified FUS binding preferentially on pre-mRNA, suggesting that these binding events occur in the nucleus^22,47–50^. Given the confirmed synaptic localization of FUS (**Fig. 1**), we wondered if a specific subset of synaptic RNAs are directly bound and regulated by FUS in these compartments. Since synapses contain few copies of different RNAs and only a small fraction of the total cellular FUS is synaptically localized, RNAs specifically bound by FUS at the synapses are likely missed in CLIP-seq datasets from total brain. Therefore, we biochemically isolated synaptoneurosomes that are enriched synaptic fractions from mouse cortex to identify synapse-specific RNA targets of FUS. Electron microscopy analysis confirmed the morphological integrity of our synaptoneurosome preparations, which contained intact pre- and postsynaptic structures (**Fig. 2a**). Immunoblot showed an enrichment of synaptic markers (PSD-95, p-CAMKII, GluN2B, GluA1, SNAP25, NXRN1), absence of nuclear proteins (Lamin B1, Histone H3) and presence of FUS in the synaptoneurosomes (**Fig. 2b** and **Supplementary 2a**). In addition, quantitative reverse transcription polymerase chain reaction (qRT-PCR) analysis showed enrichment of selected synaptic mRNAs (**Fig. 2c**).

**Fig. 2.**
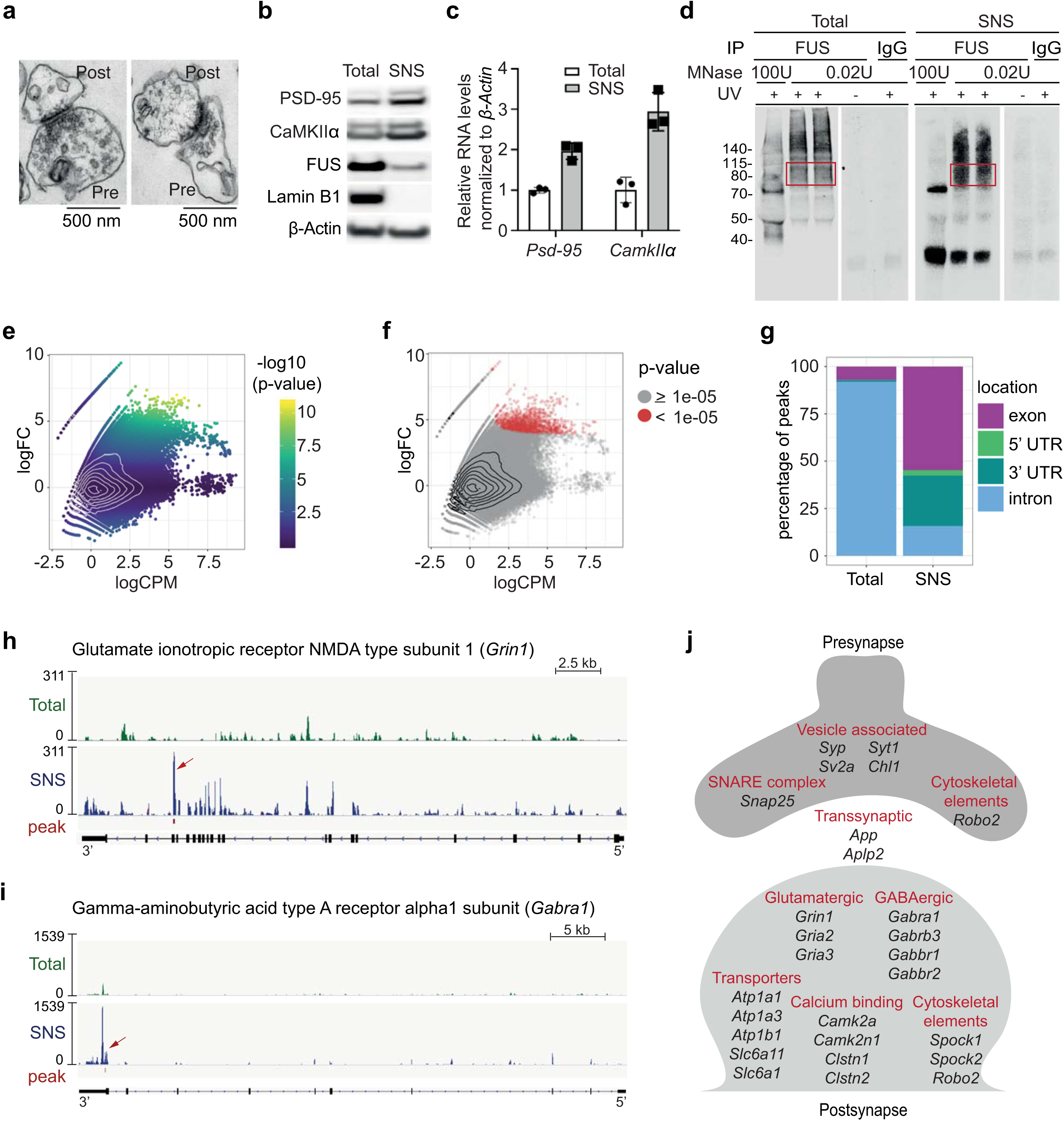
CLIP-seq on cortical synaptoneurosomes identified FUS-associated pre- and postsynaptic RNAs. **(a)** Electron microscopic images of synaptoneurosomes (SNS) from mouse cortex showing intact pre- and postsynaptic compartments. **(b)** Western blot of synaptic proteins (PSD95, p-CamKII), nuclear protein (Lamin B1) and FUS in total and SNS. **(c)** qPCR shows enrichment of PSD95, CamKII mRNAs in SNS. **(d)** Autoradiograph of FUS-RNA complexes immunoprecipitated from total homogenate and SNS and trimmed by different concentrations of micrococcal nuclease (MNase). **(e)** MA-plot of CLIPper peaks predicted in the SNS CLIP-seq sample. logCPM is the average log2CPM of each peak in the total cortex and SNS sample and logFC is the log2 fold-change between the number of reads in the SNS and total cortex sample. **(f)** Same MA-plot as E showing the selected, SNS specific peaks (p-value cutoff of 1e-05) in red. **(g)** Barplot with the percentage of SNS and total cortex specific peaks located in exons, 5’UTRs, 3’UTRs or introns. FUS binding in *Grin1* **(h)**, *Gabra1* **(i)** in total cortex (green) and SNS (blue). **(j)** Schematic with the cellular localization and function of some of the selected FUS targets.

Following a previously published method^22,51^, we used ultraviolet (UV) crosslinking on isolated synaptoneurosomes and total cortex from 1-month-old wild type mice to stabilize FUS-RNA interactions and to allow stringent immunoprecipitation of the complexes (**Supplementary Fig. 2b**). As FUS is enriched in the nucleus and only a small fraction of the protein is localized at the synapses, we prepared synaptoneurosomes from cortices of 200 mice to achieve sufficient RNA levels for CLIP-seq library preparation. The autoradiograph showed an RNA smear at the expected molecular weight of a single FUS molecule (70 kDa) and lower mobility complexes (above 115 kDa) that may correspond to RNAs bound by more than one FUS molecule or a heterogeneous protein complex (**Fig. 2d**). No complexes were immunoprecipitated in the absence of UV cross-linking or when using nonspecific IgG-coated beads. The efficiency of immunoprecipitation was confirmed by depletion of FUS in post-IP samples (**Supplementary Fig. 2c**). Finally, RNAs purified from the FUS-RNA complexes of cortical synaptoneurosomes and total cortex were sequenced and analyzed. We obtained 29,057,026 and 27,734,233 reads for the total cortex and cortical synaptoneurosome samples, respectively. 91% of the total cortex and 66% of the synaptoneurosome reads could be mapped to a unique location in the mouse reference genome (GRCm38) (**Supplementary Fig. 2d**). After removing PCR duplicates, we identified peaks using a previously published tool called CLIPper^52^, resulting in 619,728 total cortex and 408,918 synaptoneurosome peaks.

Before comparing the peaks in the two samples, we normalized the data to correct for different sequencing depths and signal-to-noise ratios^53^ (see Methods). This is especially important in our case, because the synaptoneurosome sample should contain only a subset of the FUS targets from total cortex. We wanted to filter the predicted peaks of the synaptoneurosome sample to identify genomic regions with high log2 fold-change between the synaptoneurosome and total cortex samples. Peaks with low number of reads (or no reads) in the total cortex, but high read coverage in the synaptoneurosomes correspond to regions that are putatively bound by FUS in the synapse. However, the observable number of reads per RNA in each sample strongly depends on gene expression and the number of localized RNA copies. Therefore, we did not want to use a simple read count threshold to filter and identify synapse specific peaks. Instead, we fit a count model and computed peak-specific p-values to test for differences between the synaptoneurosome and total cortex CLIP-seq enrichment (**Fig. 2e**). The normalization highlights the expected association between p-values (yellow) and log2 CPM (**Fig. 2e**).

We ranked the peaks by p-values and used a stringent cutoff of 1e-5 (**Fig. 2e**) to ensure enrichment of synaptic FUS targets. Indeed, the resulting peaks were largely devoid of intronic regions, but were enriched in exons and 3’UTRs, as was expected for synaptic FUS targets, which are mature and fully processed RNAs (**Fig. 2e** and **Supplementary Fig. 2g**). The same normalization and filtering of CLIPper peaks identified in the total cortex highlighted RNAs primarily bound by FUS in the nucleus, where the vast majority of FUS protein resides (**Supplementary Fig. 2e**). After selecting an equal number of top peaks as obtained for the synaptoneurosome sample (1560 peaks in 517 genes), corresponding to a p-value cutoff of 0.0029 (**Supplementary Fig. 2f**), we confirmed the previously reported^22^ preferential binding of FUS within intronic regions of pre-mRNAs (**Fig. 2g** and **Supplementary Fig. 2h**).

The final list of synapse-specific FUS binding sites consists of 1560 peaks in 307 RNAs (**Supplementary Table 1**), primarily localized to exons and 3’UTRs of RNAs specific to the synapses. Among those, FUS peaks on the exon of *Grin1* (Glutamate ionotropic NMDA type subunit 1) and 3’UTR of a long isoform of *Gabra1* (Gamma aminobutyric acid receptor subunit alpha-1) were exclusively detected in synaptoneurosomes, but not in total cortex (**Fig. 2h-i**). Direct binding of FUS to 3’UTR and exonic regions of its targets suggests a potential role in regulating RNA transport, local translation and/or stabilization.

### Synaptic FUS RNA targets encode essential protein components of synapse

We then wondered if the 307 synaptic FUS target RNAs were collectively highlighting any known cellular localization and function. Most RNAs are localized to either the pre- or postsynapse or they are known astrocytic markers (**Fig. 2j**). Among those are RNAs encoding essential protein members of glutamatergic (*Grin1, Gria2, Gria3*) and GABAergic synapses (*Gabra1, Gabrb3, Gabbr1, Gabbr2*), transporters, as well as components of the calcium signaling pathway, which are important for plasticity of glutamatergic synapses. An overrepresentation analysis (ORA) comparing the synaptic FUS targets to all synaptic RNAs detected in cortical mouse synaptoneurosomes by RNA-seq (logCPM >1, 1-month-old mice), revealed that FUS targets were enriched for synaptic – both pre- and postsynaptic – localization. Synaptic FUS target RNAs were enriched for gene ontology categories, such as transport, localization and trans-synaptic signaling, as well as signaling receptor binding and transmembrane transporter activity (**Supplementary Fig. 2i**).

Here we identified for the first time specific synaptic RNA targets directly bound by FUS, including those associated with glutamatergic and GABAergic networks. Our data suggests that FUS plays a critical role in maintaining synaptic integrity and organization.

### FUS binds GU-rich sequences at the synapse

While FUS has been shown to be a relatively promiscuous RNA-binding protein, preference towards GU-rich motifs has been reported in previous CLIP-seq studies^22,38,40,41^, a binding mediated via its ZnF domain^42^. To understand if FUS binding to synaptic RNA targets follows the same modalities as its nuclear targets, we explored the sequence specificity of FUS in the synapse and predicted motifs with HOMER^54^, comparing the FUS peak sequences of cortical synaptoneurosomes and total cortex samples. In accordance with previous studies, we found a degenerate GU-rich motif for intronic FUS binding sites in the total cortex (**Table 1**). The sequences of the synaptic FUS peaks in exons and 5’ UTRs revealed a “AGGUAAGU” motif which was only found in 11% and 6% of the peaks, respectively. We conclude that FUS does not have a stronger sequence preference in the synapse than in the nucleus.

**Table 1:**
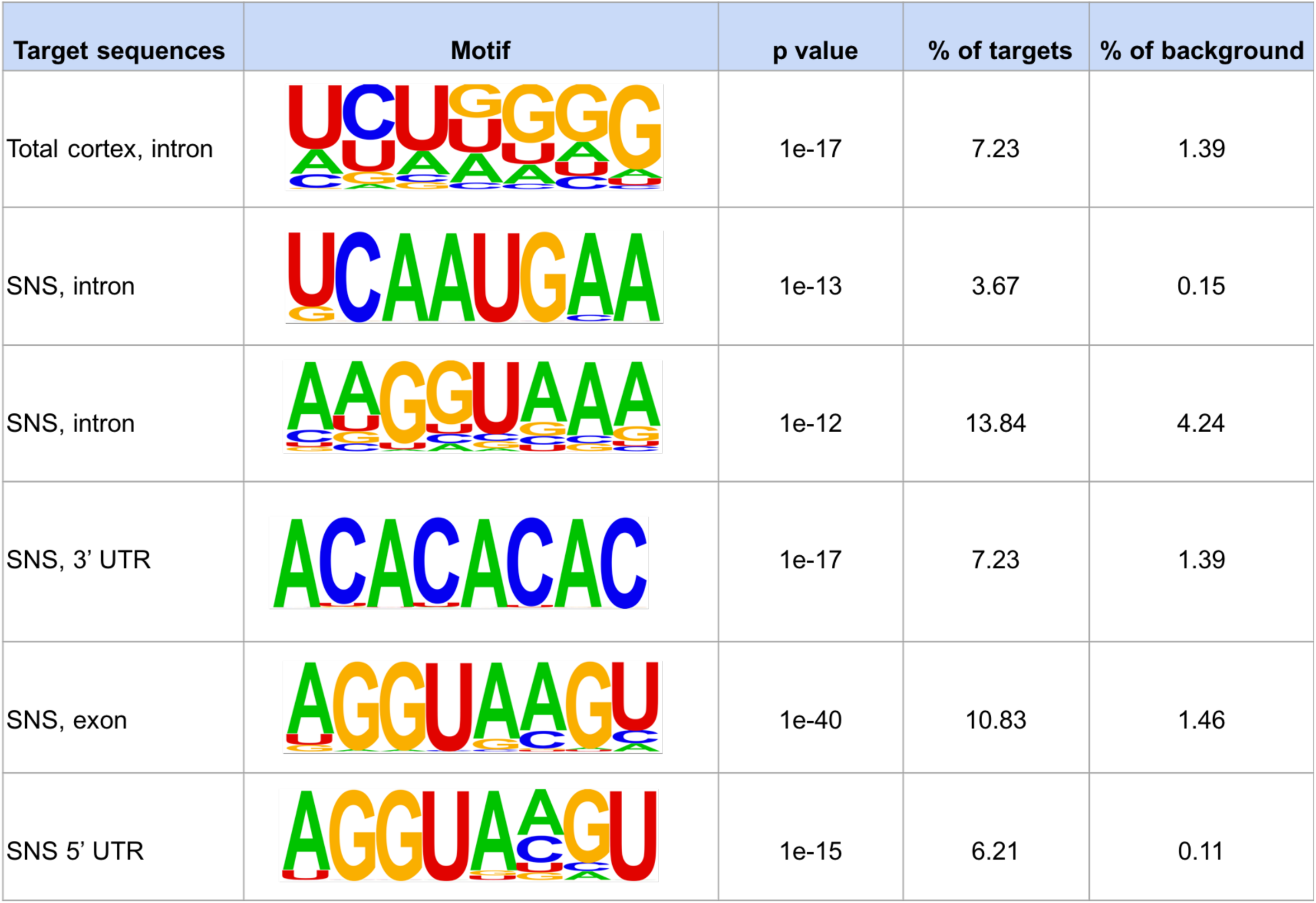
FUS binds GU-rich sequences at the synapse. Predicted sequence motifs (HOMER) in windows of size 41 centered on the position with maximum coverage in each peak. Each set of target sequences has a corresponding background set with 200,000 sequences without any CLIP-seq read coverage (they are not bound by FUS). Note: These are all motifs that were not marked as possible false positives by HOMER and that occur in more than 1% of the target sequences.

| Target sequences | Motif | p value | % of targets | % of background |
| --- | --- | --- | --- | --- |
| Total cortex, intron |  | 1e-17 | 7.23 | 1.39 |
| SNS, intron |  | 1e-13 | 3.67 | 0.15 |
| SNS, intron |  | 1e-12 | 13.84 | 4.24 |
| SNS, 3' UTR |  | 1e-17 | 7.23 | 1.39 |
| SNS, exon |  | 1e-40 | 10.83 | 1.46 |
| SNS 5' UTR |  | 1e-15 | 6.21 | 0.11 |

### Increased synaptic localization of mutant FUS protein in *Fus*^*ΔNLS/+*^ mice

In order to explore synaptic impairments associated with FUS mislocalization, we used the *Fus*^*ΔNLS*/+^ mouse model^55^. This mouse model shows partial cytoplasmic mislocalization of FUS due to a lack of the nuclear localization (NLS) in one copy of the FUS allele, closely mimicking ALS-causing mutations reported in patients. Taking advantage of two antibodies that recognize either total FUS (both full length and mutant) or only the full length protein (**Fig. 3a**), we assessed FUS protein levels in synaptoneurosomes isolated from *Fus*^*ΔNLS/+*^ mice and wild type (*Fus*^*+/+*^) of 1 and 6 months of age. We detected higher levels of total FUS in synaptoneurosomes from *Fus*^*ΔNLS/+*^ at both ages compared to *Fus*^*+/+*^ (**Fig. 3b-c, Supplementary Fig. 3a-b**). However, full length FUS levels were decreased in synaptoneurosomes of *Fus*^*ΔNLS*/+^ compared to *Fus*^*+/+*^ indicating that the truncated FUS protein is misaccumulated at the synaptic sites of *Fus*^*ΔNLS/+*^ mice.

**Fig. 3.**
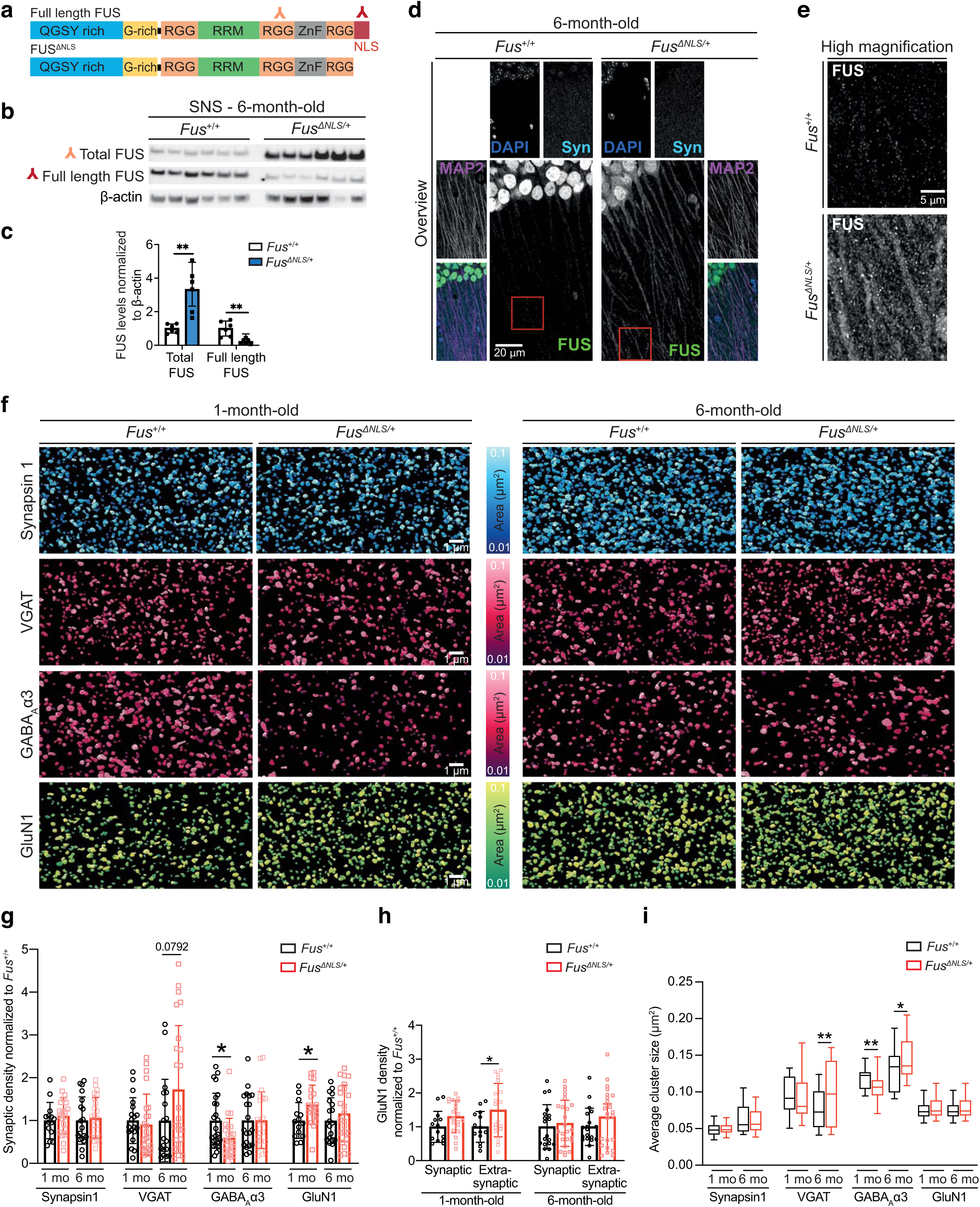
Increased synaptic FUS localization in *Fus*^*ΔNLS/+*^ mice affect GABAergic synapses. (**a**) Schematic showing specificity of antibodies used for western blot against protein domains of FUS. (**b**) Western blot of total FUS, full length FUS and actin in synaptoneurosomes isolated from *Fus*^*+/+*^ and *Fus*^*ΔNLS/+*^ mice at 6 months of age. (**c**) Quantification of total FUS and full length FUS levels in synaptoneurosomes from *Fus*^*+/+*^ and *Fus*^*ΔNLS/+*^ at 6 months of age. (**d**) Confocal images of the hippocampal CA1 area from 6-month-old mice showing higher level of FUS in the dendritic tree and synaptic compartment in *Fus*^*ΔNLS/+*^ mouse-model. On the top, low magnification pictures show the dendritic area of pyramidal cells stained with FUS (green), MAP2 (dendritic marker, magenta), Synapsin 1 (Syn, Synaptic marker, Cyan) and DAPI (Blue). Red box indicates the area imaged in the high magnification images below. (**e**) Higher magnification equivalent to the area highlighted in red in (**d**). (**f**) Representative images of staining using synaptic markers Synapsin 1, VGAT, GABA_A_α3 and GluN1 in *Fus*^*+/+*^ and *Fus*^*ΔNLS/+*^ at 1 and 6 months of age. Images were generated with Imaris and display volume view used for quantification with statistically coded surface area. Density and cluster area were analyzed. (**g**) Graph bar representation of the synaptic density of Synapsin 1, VGAT, GABA_A_α3 and GluN1 from *Fus*^*+/+*^ and *Fus*^*ΔNLS/+*^ at 1 and 6 months of age. Graph bar showing mean + SD. *p<0.05. Graphs are extracted from the same analysis shown in **Supplementary Fig. 3e-f**. The statistical analysis can be found in **Table 2.** (**h**) Colocalization analysis of GluN1 with Synapsin 1 to identify synaptic NMDAR and extrasynaptic NMDAR. Results were normalized by the control of each group. Graph bar showing mean + SD. *p<0.05. (**i**) Box and Whiskers representation of the average cluster area for each marker (Synapsin1, VGAT, GABA_A_α3 and GluN1) from 1-month and 6-month-old *Fus*^*+/+*^ and *Fus*^*ΔNLS/+*^ mice. Box showing Min to Max, *p<0.05 **p<0.01. Graphs are extracted from the same analysis shown in **Supplementary Fig. 3f-i.** The statistical analysis can be found in **Table 3.**

Confirming our biochemical evidence, immunofluorescence analyses of *Fus*^*ΔNLS*/+^ mice showed higher levels of FUS in dendritic compartments of CA1 pyramidal cells. *Fus*^*+/+*^ mice at both 1 month (**Supplementary Fig. 3c-d**) and 6 months of age (**Fig. 3d-e**) showed prominent expression of FUS in the nucleus. High magnification images highlighted the presence of FUS at the synapses, identified by co-labeling with Synapsin1. *Fus*^*ΔNLS*/+^ mice at 1 (**Supplementary Fig. 3c-d**) and 6 months of age (**Fig. 3d-e**) showed higher levels of FUS within the dendritic tree (identified with MAP2) and at the synapse compared to *Fus*^*+/+*^ mice, confirming our previous quantifications by immunoblot.

**Table 2.** Statistical analysis of synaptic density. The table reports statistical analysis of density of the synaptic markers analyzed from a minimum of 2 images from at least 4 animals per genotype (*Fus*^*+/+*^ and *Fus*^*ΔNLS/+*^) at 1 and 6 months of age. Unpaired t-test statistics, p-values, specific t-distribution (t), degrees of freedom (DF) and sample size are listed.

| unpaired t-test | 1 month |  |  | 6 months |  |  |
| --- | --- | --- | --- | --- | --- | --- |
|  | p value | t, df | sample size | p value | t, df | sample size |
| Synapsin1 | 0.4556 | t=0.7553, df=32 | +/+ =14<br>ΔNLS/+ =20 | 0.6812 | t=0.4138, df=41 | +/+ =19<br>ΔNLS/+ =24 |
| SNAP25 | 0.5320 | t=0.6319, df=32 | +/+ =14<br>ΔNLS/+ =20 | 0.085 | t=1.765, df=41 | +/+ =19<br>ΔNLS/+ =24 |
| Bassoon | 0.5821 | t=0.5567, df=28 | +/+ =18<br>ΔNLS/+ =12 | 0.4460 | t=0.7708, df=35 | +/+ =18<br>ΔNLS/+ =19 |
| VGAT | 0.6368 | t=0.4758, df=40 | +/+ =19<br>ΔNLS/+ =23 | 0.0792 | t=1.801, df=40 | +/+ =18<br>ΔNLS/+ =24 |
| GluN1 | 0.0219 | t=2.409, df=32 | +/+ =14<br>ΔNLS/+ =20 | 0.3786 | t=0.8900, df=41 | +/+ =19<br>ΔNLS/+ =24 |
| GluA1 | 0.6009 | t=0.5292, df=28 | +/+ =18<br>ΔNLS/+ =12 | 0.4885 | t=0.7000, df=35 | +/+ =18<br>ΔNLS/+ =19 |
| pCaMKII | 0.9055 | t=0.1195, df=40 | +/+ =19<br>ΔNLS/+ =23 | 0.2160 | t=1.257, df=40 | +/+ =18<br>ΔNLS/+ =24 |
| Gephyrin | 0.9878 | t=0.1531, df=88 | +/+ =43<br>ΔNLS/+ =47 | 0.5778 | t=0.5591, df=74 | +/+ =34<br>ΔNLS/+ =42 |
| GABAARα1 | 0.1368 | t=1.514, df=46 | +/+ =24<br>ΔNLS/+ =24 | 0.9611 | t=0.04906, df=44 | +/+ =20<br>ΔNLS/+ =26 |
| GABAARα3 | 0.0156 | t=2.512, df=46 | +/+ =24<br>ΔNLS/+ =24 | 0.9744 | t=0.03234, df=40 | +/+ =20<br>ΔNLS/+ =22 |

**Table 3.** Statistical analysis of synaptic cluster area. The table reports statistical analysis of area of the synaptic markers analyzed from a minimum of 2 images from at least 4 animals per genotype (*Fus*^*+/+*^ and *Fus*^*ΔNLS/+*^) at 1 and 6 months of age. Unpaired t-test statistics, p-values, specific t-distribution (t), degrees of freedom (DF) and sample size are listed.

| unpaired t-test | 1 month |  |  | 6 months |  |  |
| --- | --- | --- | --- | --- | --- | --- |
|  | p value | t, df | sample size | p value | t, df | sample size |
| Synapsin1 | 0.8249 | t=0.2214, df=363 | +/+ =14<br>ΔNLS/+ =20 | 0.643 | t=0.4639, df=393 | +/+ =19<br>ΔNLS/+ =24 |
| SNAP25 | 0.3834 | t=0.8727, df=363 | +/+ =14<br>ΔNLS/+ =20 | 0.5015 | t=0.6727, df=393 | +/+ =19<br>ΔNLS/+ =24 |
| Bassoon | 0.6022 | t=0.5217, df=363 | +/+ =18<br>ΔNLS/+ =12 | 0.7529 | t=0.315, df=393 | +/+ =18<br>ΔNLS/+ =19 |
| VGAT | 0.2819 | t=1.078, df=363 | +/+ =19<br>ΔNLS/+ =23 | 0.0028 | t=3.005, df=393 | +/+ =18<br>ΔNLS/+ =24 |
| GluN1 | 0.5437 | t=6078, df=363 | +/+ =14<br>ΔNLS/+ =20 | 0.5694 | t=0.5694, df=393 | +/+ =19<br>ΔNLS/+ =24 |
| GluA1 | 0.4303 | t=0.7896, df=363 | +/+ =18<br>ΔNLS/+ =12 | 0.4517 | t=0.7533, df=393 | +/+ =18<br>ΔNLS/+ =19 |
| pCaMKII | 0.242 | t=1.172, df=363 | +/+ =19<br>ΔNLS/+ =23 | 0.4150 | t=0.8159, df=393 | +/+ =18<br>ΔNLS/+ =24 |
| Gephyrin | 0.7467 | t=0.3233, df=363 | +/+ =43<br>ΔNLS/+ =47 | 0.2614 | t=1.125, df=393 | +/+ =34<br>ΔNLS/+ =42 |
| GABAARα1 | 0.374 | t=0.8902, df=363 | +/+ =24<br>ΔNLS/+ =24 | 0.3204 | t=0.9950, df=393 | +/+ =20<br>ΔNLS/+ =26 |
| GABAARα3 | 0.0053 | t=2.807, df=363 | +/+ =24<br>ΔNLS/+ =24 | 0.0166 | t=2.407, df=393 | +/+ =20<br>ΔNLS/+ =22 |

### Dysregulation of inhibitory synapses in *Fus*^*ΔNLS/+*^ mouse model

To explore a possible synaptic disorganization associated with mislocalization of FUS, we performed synaptic density and size analyses. Based on evidence that the hippocampal/prefrontal cortex connectome participates in memory encoding and recalling^56^ and that CA1 hippocampal excitatory and inhibitory synapses are highly similar to the cortical synapses^57–60^, we explored the possible synaptic changes triggered by FUS mislocalization in the CA1 hippocampal region. We analyzed both *Fus*^*+/+*^ and *Fus*^*ΔNLS*/+^ mice, using presynaptic and postsynaptic markers. Density and area analyses were performed as shown in **Supplementary Fig. 3e**. At the presynapse, we quantified the density of the SNARE associated protein SNAP25^61^ (synaptic RNA target of FUS) and the presynaptic active zone marker Bassoon^45^. The density of inhibitory synapses was assessed using VGAT^62^ (presynaptic). At the postsynapse, we quantified the density of postsynaptic glutamatergic receptor GluN1^63^ (synaptic RNA target of FUS and obligatory subunit of all NMDAR) and GluA1^64^ (obligatory subunit of AMPAR), as well as postsynaptic GABAergic receptors containing α1 subunit (GABA_A_α1; synaptic RNA target of FUS) and α3 (GABA_A_α3)^65^. We also assessed the number of active excitatory synapses using phospho-CaMKII (pCaMKII) as well as functional inhibitory synapses using Gephyrin^66^.

At 1 month of age in *Fus*^*ΔNLS*/+^ mice, we did not observe significant changes at the presynaptic site, suggesting a normal axonal and axon terminal development and functions. However, at the postsynaptic sites, we observed a significant increase of NMDAR (p=0.0219) and a significant decrease of GABA_A_α3 receptors (p=0.0156) (**Fig. 3f-g, Supplementary Fig. 3f** and **Table 2**). Moreover at 1 month of age, *Fus*^*ΔNLS*/+^ mice showed significantly more NMDAR located at the extrasynaptic site (p=0.0433) (**Fig. 3h**). Interestingly, the size of the GABA_A_α3 clusters was significantly decreased in *Fus*^*ΔNLS*/+^ mice (p=0.0053) at 1 month of age (**Fig. 3f, i, Supplementary Fig. 3h** and **Table 3**). We did not record changes in the number of Synapsin1, Bassoon, SNAP25, VGAT, GluA1, GABA_A_α1, Gephyrin or pCaMKII, suggesting either an increase of silent synapses, immature synapses or an increase of the number of NMDAR in the dendritic shaft together with a decrease of GABA_A_α3 synaptic clustering. These results suggested a hyperexcitability profile during developmental stages.

At 6 months of age, we did not observe significant changes in the density of pre or postsynaptic markers (**Fig. 3f-g** a**nd Supplementary Fig. 3g**), suggesting a normal maturation of the synaptic network despite developmental synaptic dysregulation described above. However, SNAP25 (p=0.085) and VGAT (p=0.0792) trended towards an increased density, suggesting a potential alteration at inhibitory presynaptic sites (**Supplementary Fig. 3g and Table 2**). This interpretation was confirmed by an increase of the area of the presynaptic marker VGAT (p=0.0028) and of the size of GABA_A_α3 clusters at the postsynaptic site (p=0.0166) (**Fig. 3i, Supplementary Fig. 3i** and **Table 3**), while GluN1 clusters appeared unaffected. Increase in VGAT suggested an elevated number of presynaptic GABAergic vesicles, which was confirmed by EM analyses in older mice (Scekic-Zahirovic, Sanjuan-Ruiz et al., co-submitted manuscript). Correlatively, increase of GABA_A_α3 cluster size suggested an increase in the trafficking of GABA_A_R at the postsynaptic site. This occurred, however, without an increase of the anchoring protein Gephyrin, suggesting instable structure of the inhibitory postsynaptic sites. Altogether, our results show alterations of both glutamatergic and GABAergic synapses during developmental synaptogenesis (1 month of age), while only GABAergic synapses appeared affected at a later time point (6 months of age). This suggests a potential role for FUS in synaptogenesis and network wiring and synaptic maintenance, with a selective exacerbation of inhibitory synaptic defects with age.

### *Fus*^*ΔNLS/+*^ mice show age-dependent synaptic RNA alterations

FUS plays an essential role in RNA stabilization^23,24^ and transport^20^. Therefore, we used RNA-seq to investigate the consequences of increased synaptic levels of mutated FUS in *Fus*^*ΔNLS*/+^ mice (**Fig. 4a**). We isolated RNA from six biological replicates of synaptoneurosomes and paired total cortex samples from *Fus*^*+/+*^ and *Fus*^*ΔNLS*/+^ mice at 1 and 6 months of age and prepared poly-A-selected libraries for high-throughput sequencing. As a control, we also sequenced the nuclear fraction from 4 biological replicates of *Fus*^*+/+*^ mice at 1 month of age. For quality control, we computed principal components of all samples and all expressed genes (see methods for details) and found a clustering by sample condition and age (**Supplementary Fig. 4b-c).**

**Fig. 4.**
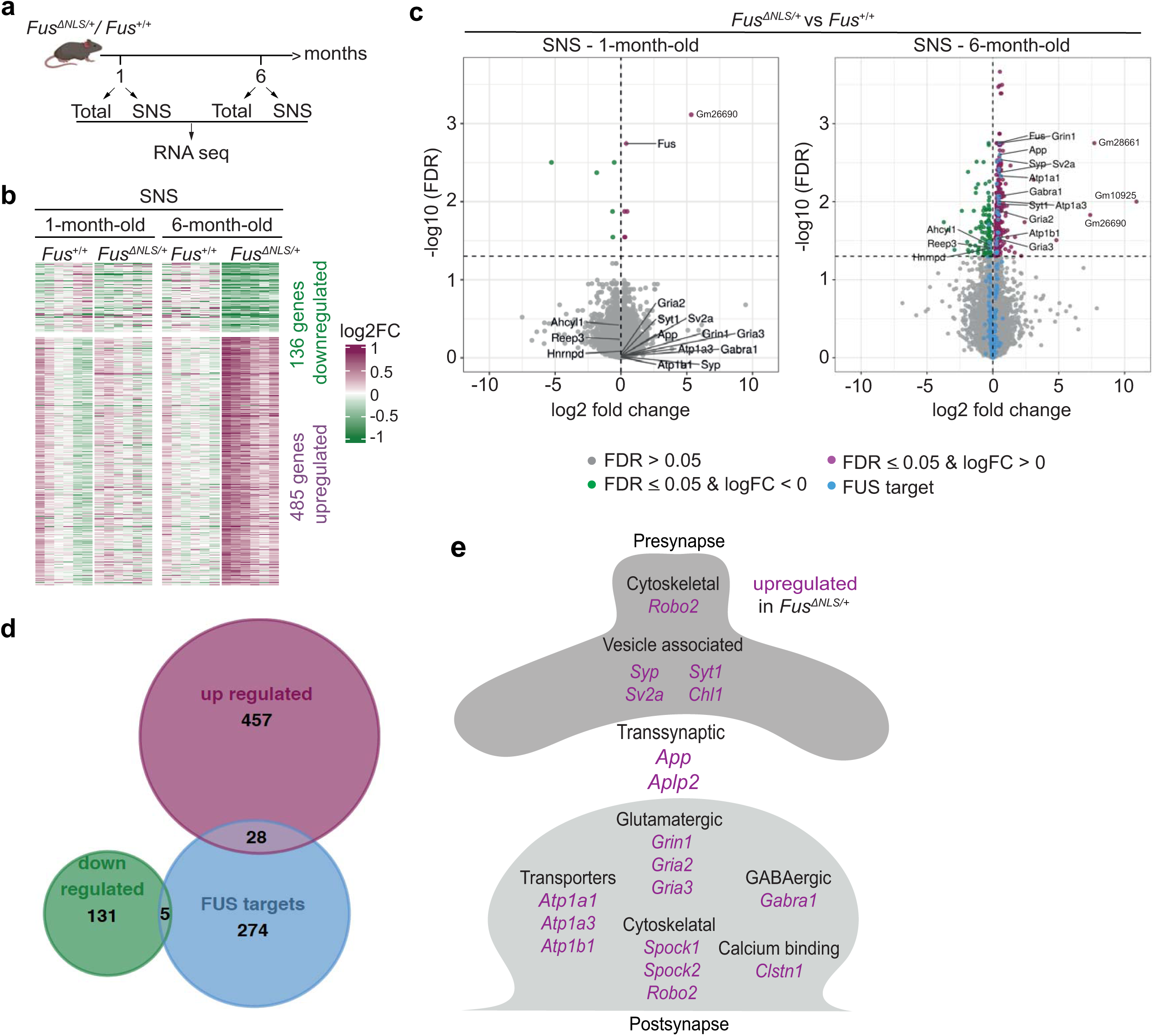
Age-dependent alterations in the synaptic RNA profile of *Fus*^*ΔNLS/+*^ mouse cortex. (**a**) Outline of the RNA-seq experiment. (**b**) Heatmap from the set of up- and downregulated genes in SNS of *Fus*^*ΔNLS/+*^ at 6-months compared to *Fus*^*+/+*^. Genes are on the rows and the different samples on the columns. The color scale indicates the log2FC between the CPM of each sample and mean CPM of the corresponding *Fus*^*+/+*^ samples at each time point [sample logCPM – mean (logCPM of *Fus*^*+/+*^ samples)]. (**c**) Volcano plots showing the log2 fold change of each gene and the corresponding minus log10 (FDR) of the differential gene expression analysis comparing *Fus*^*ΔNLS/+*^ SNS to *Fus*^*+/+*^ SNS at 1 month (left panel) and 6 months of age (right panel). The horizontal line marks the significance threshold of 0.05. Significantly downregulated genes are highlighted in green, upregulated genes in purple and all FUS targets identified in the CLIP-seq data in blue. (**d**) Venn diagram of the sets of significantly up- and downregulated genes (SNS of *Fus*^*ΔNLS/+*^ vs. *Fus*^*+/+*^ at 6 months of age) and the SNS FUS target genes identified by our FUS CLIP-seq. (**e**) Schematic of the cellular localization of the differentially expressed FUS targets in SNS of *Fus*^*ΔNLS/+*^ mice at 6 months of age.

We compared the expressed genes in our synaptoneurosomes (15087 genes) with the forebrain synaptic transcriptome^67^ (14073 genes) and the vast majority of detected RNAs (13475) were identical between the two studies (**Supplementary Fig. 4a**). The small differences in the two transcriptomes can be explained by differences in the used synaptoneurosome protocols and the brain region (frontal cortex versus forebrain).

We conducted four differential gene expression analyses, comparing *Fus*^*ΔNLS/+*^ to *Fus*^*+/+*^ replicates separately for the total cortex and synaptoneurosomes at both time points (for full lists see **Supplementary Tables 2-5**). A false discovery rate (FDR) cutoff of 0.05 was used to define significant differential expression. Only three and five RNAs were differentially expressed (DE) in the *Fus*^*ΔNLS/+*^ samples of the total cortex at 1 and 6 months of age, respectively (**Supplementary Fig. 4f and Supplementary Tables 2-3**). However, in the synaptoneurosomes, we identified 11 and 594 RNAs differentially abundant at 1 and 6 months, respectively (**Supplementary Tables 4-5**). 136 RNAs were decreased and 485 RNAs were increased in the *Fus*^*ΔNLS/+*^ mice at 6 months of age compared to synaptoneurosomes from *Fus*^*+/+*^ mice (**Fig. 4b**). The significantly increased RNAs in *Fus*^*ΔNLS/+*^ mice at 6 months were enriched in gene ontology (GO) categories such as synaptic signaling, intrinsic component of membrane and transporter activity (**Supplementary Fig. 4d**), while those that were decreased in abundance were associated with cytoskeletal organization and RNA metabolism (**Supplementary Fig. 4e**).

At 6 months of age, the log2 fold changes of the altered RNAs are consistently negative or positive in all *Fus*^*ΔNLS/+*^ synaptoneurosome replicates (**Fig. 4c**). At 1 month of age, the log2 fold changes of the *Fus*^*ΔNLS/+*^ synaptoneurosome replicates are mostly neutral (white color on the heatmap) indicating that alterations in RNA abundance are age-dependent and not detectable as early as 1 month of age. In the total cortical samples at 6 months of age, some of the replicates show a similar trend as the synaptoneurosome samples, but it seems that the effects cannot be detected because synaptic RNAs are too diluted (**Supplementary Fig. 4g**). Overall, we found synapse-specific differential RNA abundance at 6 months in the *Fus*^*ΔNLS/+*^ mice, but not in the total cortex.

While most of the 594 differentially abundant RNAs (**Supplementary Table 5**) were not direct FUS targets, 33 altered RNAs are synaptic targets of FUS. The altered synaptic transcriptome, along with the impaired expression of a subset of FUS RNA targets in *Fus*^*ΔNLS/+*^ mice, suggests direct and indirect effects of mutant FUS at the synapses (**Fig. 4d**). FUS targets with known synaptic functions that are altered in *Fus*^*ΔNLS/+*^ are represented in **Fig. 4e**. Most of those RNAs show exonic FUS binding on our CLIP-seq analysis (**Supplementary Fig. 5-6, Supplementary Table 1**), with the exception of *Gria 3, Spock1, Spock2* (**Supplementary Fig. 6b, f-g**) and *Gabra1* (**Supplementary Fig. 7**), which are bound by FUS at their 3’UTR. Altered FUS targets include RNAs encoding presynaptic vesicle associated proteins, transsynaptic proteins, membrane proteins, receptors associated with glutamatergic and GABAergic pathways. Our results suggest that mislocalization of FUS leads to mild alterations in the synaptic RNA profile that may affect synaptic signaling and plasticity. Our data indicate that synaptic RNA alterations may occur at an asymptomatic age and represent one of the early events in disease pathogenesis.

## Discussion

In this study, we identified for the first-time synaptic RNA targets of FUS combining cortical synaptoneurosome preparations with CLIP-seq. Additionally, synaptic RNA levels were found to be altered in a *Fus*^*ΔNLS/+*^ mouse model at 6 months of age. Along with these results, we assessed FUS localization at the synaptic site using a combination of super-resolution microscopy approaches. Altogether, our results point to a critical role for FUS at the synapse and indicate that increased synaptic FUS localization at presymptomatic stages of ALS-FUS mice triggers early alterations of synaptic RNA content and misregulation of the GABAergic network. These early synaptic changes mechanistically explain the behavioral dysfunctions that these mice develop (Scekic-Zahirovic, Sanjuan-Ruiz et al., co-submitted manuscript).

RNA transport and local translation ensure fast responses with locally synthesized proteins essential for plasticity^21,68,69^. CLIP-seq using synaptoneurosome preparations from mouse cortex demonstrated that FUS not only binds nuclear RNAs, but also those that are localized at the synapses. Both pre- and postsynaptic localization of the identified targets correlated with the subcellular localization of FUS in both synaptic compartments. Moreover, by CLIP-seq on synaptoneurosomes, we identified that FUS binds RNAs encoding GABA receptor subunits (*Gabra1, Gabrb3, Gabbr1, Gabbr2*) and glutamatergic receptors (*Gria2, Gria3, Grin1*) previously known to be localized at dendritic neuropils^70^. FUS binding on synaptic RNAs is enriched on 3’UTRs and/or exonic regions, as revealed by our synaptoneurosome CLIP-seq dataset, suggesting that FUS might play a role in regulating local translation or transport of these targets.

Synaptic analyses at presymptomatic ages of *Fus*^*ΔNLS/+*^ mice revealed interesting changes. Our results showed a major effect on inhibitory synapses at 1 and 6 months of age. We explored GABA_A_R density and found changes in α3-containing GABA_A_R. GABA_A_α3 is expressed at the postsynaptic site of monoaminergic synapses^71^, and have been shown to be involved in fear and anxiety behavior, and mutations in the *Gabra3* subunit resulted in an absence of inhibition behavior^72–74^. Changes in GABA_A_α3 and not GABA_A_α1-containing receptor suggested that only monoaminergic neurons were affected in the *Fus*^*ΔNLS/+*^ mouse model. These results are well aligned with a contemporaneous study (Scekic-Zahirovic, Sanjuan-Ruiz et al., co-submitted manuscript), which showed specific behavioral changes that can be linked to monoaminergic networks. Interestingly at 1 month of age, *Fus*^*ΔNLS/+*^ mice showed an increase of NMDAR associated with a decrease in GABA_A_α3. These results suggested a role for FUS during synaptogenesis in regulating postsynaptic receptor composition as previously suggested^23,28,75^. In 1-month-old *Fus*^*ΔNLS/+*^ mice, NMDARs were enriched at the extrasynaptic sites, which, together with the decrease in GABA_A_α3, suggested an hyperexcitability profile during development. We hypothesize that abnormal activity during developmental stages could result in abnormal network connection. *Fus*^*ΔNLS/+*^ mice at 6 months of age showed higher density of presynaptic inhibitory boutons, pointing toward a compensatory mechanism at the GABAergic synapses to overcome the hyperexcitability profile observed during development. Moreover at 6 months of age, *Fus*^*ΔNLS/+*^ mice also displayed higher density of SNAP25, present at both inhibitory and excitatory synapses^61,76^, but we did not explore if this increase was specific for the GABAergic network.

Interestingly, the cluster size of VGAT, which is involved in the transport of GABA in the presynaptic vesicles^77^, was increased in *Fus*^*ΔNLS/+*^ mice at 6 months of age. Increase of the cluster size would suggest that either more vesicles were present at the presynapse, or an increase of VGAT protein per vesicle. We also observed an increase in GABA_A_α3 cluster size and their density in 6-month-old *Fus*^*ΔNLS/+*^ mice. Surprisingly, we did not observe an increase in Gephyrin, a postsynaptic protein responsible for anchoring GABAR at the postsynaptic site^78,79^. Gephyrin interacts at the postsynaptic site with GABAR at a ratio 1:1^80^, suggesting that inhibitory synapses in the *Fus*^*ΔNLS/+*^ model were unstable at 6 months of age with an excess of GABAR poorly anchored at the postsynaptic site, which could lead to malfunction of the inhibitory network. In correlation, *Fus*^*ΔNLS*/+^ mice showed behavioral changes overtime with disinhibition and hyperactivity behaviors as early as 4 months of age, associated with a decrease in the number of inhibitory neurons at 22-month-old (Scekic-Zahirovic, Sanjuan-Ruiz et al., co-submitted manuscript). Altogether, these results suggest that increased level of extranuclear FUS during development led to abnormal synaptogenesis affecting the GABAergic system over time.

Using the *Fus*^*ΔNLS*/+^ mouse model, we found that accumulation of mislocalized mutant FUS at the synapses altered the synaptic RNA content as early at 6 months of age. These alterations include FUS target RNAs that are associated with glutamatergic (*Grin1, Gria2, Gria3*) and GABAergic (*Gabra1*) synapses. These targets were found with increased synaptic localization in *Fus*^*ΔNLS/+*^. An impairment of genes associated with the GABAergic network in the frontal cortex of both young (5-month-old) and old (22-month-old) *Fus*^*ΔNLS*/+^ mice has been shown by an independent study (Scekic-Zahirovic, Sanjuan-Ruiz et al., co-submitted manuscript). Importantly, this ALS-FUS mouse model developed behavioral deficits, including hyperactivity and social disinhibition, suggesting defects in cortical inhibition. Our data supports that phenotypic manifestations in *Fus*^*ΔNLS*/+^ mice could be due to synaptic RNA alterations caused by mutant FUS at synapses. Moreover, mutant FUS-associated synaptic RNA alterations precede in ALS-FUS mice as suggested in our data. However, the precise mechanism of how FUS regulates these targets is yet to be determined.

CLIP-seq from synaptoneurosomes showed that FUS binds selectively to specific GABA receptor subunits encoding mRNAs: *Gabra1, Gabrb3, Gabbr1, Gabbr2*. Other RNA-binding proteins, such as fragile X mental retardation protein (FMRP), Pumilio 1, 2 and cytoplasmic polyadenylation binding element binding protein (CPEB) have also been shown to bind GABAR subunit mRNAs by CLIP-seq^81^. Whether all these proteins act in concert to locally regulate the expression of GABAR subunits at synapses needs to be investigated. Interestingly, FUS interacts with FMRP, a well-studied protein known to regulate local translation^82^. Long 3’ UTRs have been suggested to promote increased binding of RBPs and miRNAs which control the translation of these mRNAs^83^. Our CLIP-seq from synaptoneurosomes showed that FUS binds to the long 3’ UTR containing isoform of *Gabra1* (**Supplementary Fig. 7**) indicating that FUS may be directly involved in regulating the protein expression of *Gabra1* at the synapses. Furthermore, we found increased levels of *Gabra1* mRNA in synaptoneurosome preparations from *Fus*^*ΔNLS*/+^ mice. It is important to study whether elevated levels of FUS at the synapse may directly impact *Gabra1* levels via mRNA stabilization or local translation leading to altered regulation of inhibitory network. Overall, our findings highlight the role of FUS in synaptic RNA homeostasis possibly through regulating RNA transport, RNA stabilization and local translation.

## Materials and Methods

### Experimental models

Mice housing and breeding were in accordance with the Swiss Animal Welfare Law and in compliance with the regulations of the Cantonal Veterinary Office, Zurich. We used 1- to 6-month-old C57/Bl6 mice or *Fus*^*+/+*^/*Fus*^*ΔNLS/+*^ mice with genetic background (C57/Bl6). Wild type and heterozygous *Fus*^*ΔNLS/+*^ mice with genetic background (C57/Bl6)^55^ were bred and housed in the animal facility of the University of Zurich.

### Immunofluorescence staining for brain sections

Mice were anesthetized by CO_2_ inhalation before perfusion with PBS containing 4% paraformaldehyde and 4% sucrose. Brains were harvested and post-fixed overnight in the same fixative and then stored at 4°C in PBS containing 30% sucrose. Sixty μm-thick coronal sections were cut on a cryostat and processed for free-floating immunofluorescence staining. Brain sections were incubated with the indicated primary antibodies for 48 h at 4°C followed by secondary antibodies for 24h at 4°C. The antibodies were diluted in 1X Tris Buffer Saline solution containing 10% donkey serum, 3% BSA, and 0.25% Triton-X100. Sections were then mounted on slides with Prolong Diamond (Life Technologies) before confocal microscopy.

### STED super-resolution imaging and analysis

Super-resolution STED (Stimulated emission depletion microscopy) images of FUS and synaptic markers were acquired on a Leica SP8 3D, 3-color gated STED laser scanning confocal microscope. Images were acquired in the retrospenial cortical area in the layer 5 and in the molecular layer of the hippocampal CA1 area. A 775 nm depletion laser was used to deplete both 647 and 594 dyes. The powers used for depletion lasers, the excitation laser parameters, and the gating parameters necessary to obtain STED resolution were assessed for each marker. 1 μm-thick Z-stacks of 1024 × 1024-pixel images at 40 nm step size were acquired at 1800 kHz bidirectional scan rate with a line averaging of 32 and 3 frame accumulation, using a 100X (1.45) objective with a digital zoom factor of 7.5, yielding 15.15 nm pixels resolution.

STED microscopy data were quantified from at least 2 image stacks acquired from 2 *Fus*^*+/+*^ adult mice. The STED images were deconvolved using Huygens Professional software (Scientific Volume Imaging). Images were subsequently analyzed using Imaris software. Volumes for each marker were generated using smooth surfaces with details set up at 0.01 m. The diameter of the largest sphere was set up at 1 μm. Threshold background subtraction methods were used to create the surface, and the threshold was calculated for each marker and kept constant. Surfaces were then filtered by setting up the number of voxels >10 and <2000 pixels. Closest neighbor distance was calculated using integrated distance transformation tool in Imaris. Distances were then organized and statistically analyzed using mean comparison and t-test comparison. Distances greater than 200 nm were removed from the analysis, and average distance were analyzed.

### Neuronal primary cultures

Primary neuronal cell cultures were prepared from postnatal (P0) pups. Briefly, hippocampus and cortex were isolated. Hippocampi were treated with trypsin (0.5% w/v) in HBSS-Glucose (D-Glucose, 0.65 mg/ml) and triturated with glass pipettes to dissociate tissue in Neurobasal medium (NB) supplemented with glutamine (2 mM), 2% B27, 2.5% Horse Serum, 100U penicillin-streptomycin and D-Glucose (0.65 mg/ml). Hippocampal cells were then plated onto poly-D-lysine coated 18×18 mm coverslips (REF) at 6 × 10^4^ cells/cm^2^ for imaging, and for biochemistry at high density (8 x 10^4^ cells/cm^2^). Cells were subsequently cultured in supplemented Neurobasal (NB) medium at 37°C under 5% CO_2_, one-half of the medium changed every 5 days, and used after 15 days in vitro (DIV). Cortex were dissociated and plated similarly to hippocampal cells in NB supplemented with 2% B27, 5% horse serum, 1% N2, 1% glutamax, 100U penicillin-streptomycin and D-Glucose (0.65 mg/ml).

### Direct Stochastic Optical Reconstruction Microscopy (dSTORM)

Super-resolution images were acquired on a Leica SR Ground State Depletion 3D / 3 color TIRFM microscope with an Andor iXon Ultra 897 EMCCD camera (Andor Technology PLC). DIV15-18 mouse primary neurons were fixed for 20 min in 4% PFA – 4% sucrose in PBS. Primary antibodies were incubated overnight at 4% in PBS containing 10% donkey serum, 3% BSA, and 0.25% Triton X-100. Secondary antibodies were incubated at RT for 3 h in the same buffer. After 3 washes in PBS, the cells were re-fixed with 4%PFA for 5 min. The coverslips were then washed over a period of 2 days at 4°C in PBS to remove non-specific binding of the secondary antibodies. Coverslips were mounted temporarily in an oxygen scavenger buffer (200mM phosphate buffer, 40% glucose, 1M cysteamine hydrochloride (M6500 Sigma), 0.5mg/mL Glucose-oxydase, 40ug/mL Catalase) to limit oxidation of the fluorophores during image acquisition. The areas of capture were blindly selected by direct observation in DIC. Images were acquired using a 160X (NA 1.43) objective in the TIRF mode North direction with a penetration of 200 nm. Far red channels (Alexa 647 or 660) were acquired using a 642 nm laser. Red channels (Alexa 568 or 555) were acquired using a 532 nm laser. Green channel (Alexa 488) was acquired using 488 nm laser. Images were acquired in 2D. The irradiation intensity was adjusted until the single molecule detection reached a frame correlation <0.25. Detection particle threshold was defined between 20-60 depending on the marker and adjusted to obtain a number of events per frame between 0 and 25. The exposure was maintained at 7.07 ms and the EM gain was set at 300. The power of depletion and acquisition was defined for each marker and kept constant during acquisition. The number of particles collected were maintained constant per markers and between experiments. At least 3 independent cultures or coverslips were imaged per marker.

### Super-resolution image processing and analysis

Raw GSD images were processed using a custom-made macro in Fiji to remove background by subtraction of a running median of frames (300 renewed every 300 frames) and subtracting the previously processed image once background was removed^84^. A blur (0.7-pixel radius) per slice prior to median subtraction was applied to reduce the noise further. These images were then processed using Thunderstorm plugin in Imagej. Image filtering was performed using Wavelet filter (B-spline, order 3/scale2.0). The molecules were localized using centroid of connected components, and the peak intensity threshold was determined per marker/dye to maintain an XY uncertainty <50. Sub-pixel localization of molecules was performed using PSF elliptical gaussian and least squared fitting methods with a fitting radius of 5 pixels and initial sigma of 1.6 pixels. Images were analyzed using Bitplane Imaris software v.9.3.0 (Andor Technology PLC). Volumes for each marker were generated using smooth surfaces with details set up at 0.005. The diameter of the largest sphere was set up at 1 μm. A threshold background subtraction method was used to create the surface and threshold was calculated and applied to all the images of the same experiment. Surfaces were then filtered by setting up the area between 0.01-1 μm^2^. The closest neighbor distance was processed using the integrated distance transformation tool in Imaris. Distances were then organized and statistically analyzed using median comparison and ANOVA and Fisher’s Least Significant Difference (LSD) test. Distances greater than 100 nm were removed from the analysis, and average distance were analyzed.

### Preparation of synaptoneurosomes from mouse brain tissues

Synaptoneurosomes were prepared based on previously published protocols^85,86^ with slight modifications. The freshly harvested cortex tissue homogenized using dounce homogenizer for 12 strokes at 4°C in buffer (10%w/v) containing pH 7.4, 10 mM 4-(2 hydroxyethyl)-1-piperazineethanesulfonic acid (HEPES; Biosolve 08042359), 0.35 M Sucrose, 1 mM ethylenediaminetetraacetic acid (EDTA; VWR 0105), 0.25 mM dithiothreitol (Thermo Fisher Scientific R0861), 30 U/ml RNAse inhibitor (Life Technologies N8080119) and complete-EDTA free protease inhibitor cocktail (Roche 11836170001, PhosSTOP (Roche 04906845001). 200ul of the total homogenate were saved for RNA extraction or western blot analysis. The remaining homogenate was spun at 1000g, 15 min at 4°C to remove the nuclear and cell debris. The supernatant was sequentially passed through three 100 μm nylon net filters (Millipore NY1H02500), followed by one 5 μm filter (Millipore SMWP013000). The filtrate was resuspended in 3 volumes of SNS buffer without sucrose and spun at 2000g, 15 min at 4°C to collect the pellet containing synaptoneurosomes. The pellets were resuspended in RIPA buffer for western blot or in qiazol reagent for RNA extraction.

### Cross-Linking Immunoprecipitation and high-throughput sequencing (CLIP-seq)

Total lysate and synaptoneurosomes isolated from cortex tissue of 1-month-old C57Bl/6 mice were UV crosslinked (100 mJ/cm^2^ for 2 cycles) using UV Stratalinker 2400 (Stratagene) and stored at −80°C until use. For the total sample, cortex tissue was dissociated using a cell strainer of pore size 100 μm before crosslinking. We used cortex from 200 mice to prepare SNS and two mice for the total cortex sample. We used a mouse monoclonal antibody specific for the C-terminus of FUS (Santa Cruz) to pull down FUS associated RNAs using magnetic beads. After immunoprecipitation, FUS-RNA complexes were treated with MNAse in mild conditions and the 5’ end of RNAs were radiolabeled with P^32^-gamma ATP. Samples run on SDS-gel (10% Bis Tris) were transferred to nitrocellulose membrane and visualized using FLA phosphorimager. RNAs corresponding to FUS-RNA complexes were purified from the nitrocellulose membrane and strand-specific paired-end CLIP libraries were sequenced on HiSeq 2500 for 15 cycles.

### Bioinformatic analysis of CLIP-seq data and identification of FUS targets

Low quality reads were filtered and adapter sequences were removed with Trim Galore! (Krueger, F., TrimGalore. Retrieved February 24, 2010, from https://github.com/FelixKrueger/TrimGalore). Reads were aligned to the mouse reference genome (build GRCm38) using STAR version 2.4.2a^87^ and Ensembl gene annotations (version 90). We allowed a maximum of two mismatches per read (--outFilterMismatchNmax 2) and removed all multimapping reads (--outFilterMultimapNmax 1). PCR duplicates were removed with Picard tools version 2.18.4 (“Picard Toolkit.” 2019. Broad Institute, GitHub Repository. http://broadinstitute.github.io/picard/; Broad Institute). Peaks were called separately on each sample with CLIPper^52^ using default parameters.

To identify regions that are specifically bound by FUS in the SNS sample but not the total cortex sample, we filtered the peaks based on an MA plot. For each peak, we counted the number of overlapping reads in the SNS (x) and total cortex samples (y). M (log2 fold change) and A (average log2 counts) were calculated as follows: 

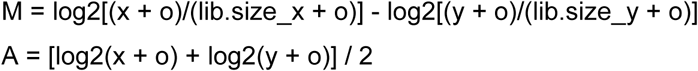

where o = 1 is an offset to prevent a division by 0 and lib.size_x and lib.size_y is the effective library size of the two samples: the library size (number of reads mapping to the peaks) multiplied by the normalization factor obtained from “calcNormFactors” using the trimmed mean of M-values^88^ method. The M and A values of all CLIPper peaks identified in the SNS sample were plotted against each other (x-axis A, y-axis M). The plot was not centered at a log2FC of 0. Therefore, we fitted a LOESS (locally estimated scatterplot smoothing) curve for normalization (loess (formula=M∼A, span=1/4, family=“symmetric”, degree=1, iterations=4)). We computed the predicted M values (fitted) for each A value and adjusted the M values by the fit (adjusted M = M – fitted M). After adjustment, the fitted LOESS line crosses the y-axis at 0 with slope = 0 in the adjusted MA-plot.

For ranking purposes, we computed p-values for each peak with the Bioconductor edgeR package^88^. We computed the common dispersion of the peaks at the center of the main point cloud (−3 < y < 1 in raw MA-plot) and not the tagwise dispersion because we are lacking replicate information. Peak specific offsets were computed as log (lib.size*norm.factors) where norm.factors are the normalization factors. The fitted M-values were subtracted from the peak specific offsets to use the adjustments from the LOESS fit for the statistical inference. We fit a negative binomial generalized linear model to the peak specific read counts using the adjusted offsets. We want to test for differential read counts between the synaptoneurosome and total cortex sample (∼group). A likelihood ratio test^89^ was run on each peak to test for synaptoneurosome versus total cortex differences.

We compared the sets of peaks obtained from different p-value cutoffs (Supplementary **Fig. 2g**) and choose the most stringed cutoff of 1e-5 because it showed the strongest depletion of intronic peaks and strongest enrichment of exonic and 3’UTR peaks. CLIPper annotated each peak to a gene and we manually inspected the assigned genes and removed wrong assignments caused by overlapping gene annotations.

Total cortex-specific peaks (regions that are exclusively bound in the total cortex sample but not the SNS sample) were computed with the same approach: the M values were computed as 

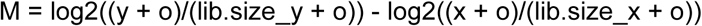

and we used a p-value cutoff of 0.0029825 because that resulted in an identical number of SNS-specific peaks.

For the over representation analysis (ORA) we applied the “goana” function from the limma R package using the gene length as covariate^90^. As background set, we used all genes with a cpm of at least 1 in all RNA-seq samples of synaptoneurosomes from 1-month-old mice.

RNA motifs of length 2-8 were predicted with HOMER^54^. To help with the motif finding, we decided to use input sequences of equal length because the lengths of the predicted peaks varied a lot. We define the peak center as the median position with maximum read coverage. Then, we centered a window of size 41 on the peak center of each selected peak and extracted the genomic sequence. We generated background sequences for each set of target sequences. A background set consists of 200,000 sequences of length 41 from random locations with the same annotation as the corresponding target set (intron, exon, 3’ UTR or 5’ UTR). All background sequences are from regions without any read coverage in the corresponding CLIP-seq sample to ensure that the background sequences are not bound by FUS.

### RNA extraction and high-throughput sequencing (RNA-seq)

Cortex tissue was isolated from 1 and/or 6-month-old *Fus*^*ΔNLS*/+^ and *Fus*^*+*/+^ mice. Paired total cortex (200 μl) and SNS sample was obtained from a single mouse per condition using filtration protocol as previously described. Briefly, frozen total and SNS samples were mixed with Qiazol reagent following the manufacturer’s recommendations and incubated at RT for 5 min. Two hundred microliters of chloroform were added to the samples and mixed for 15s and then centrifuged for 15 min (12,000g, 4°C). To the upper aqueous phase collected, five hundred microliters of isopropanol and 0.8 μl of glycogen was added and incubated at RT for 15 minutes. The samples were centrifuged at 10,000 rpm for 10 min. After centrifugation at 12,000g for 15 min, the isopropanol was removed and the pellet was washed with 1 ml of 70% ethanol and samples were centrifuged for 5 min at 7500g. Ethanol was discarded and the RNA pellet was air-dried and dissolved in nuclease free water and further purified using the RNeasy Mini Kit including the DNAse I digestion step. The concentration and the RIN values were determined by Bioanalyzer. 150 ng of total RNA were used for Poly A library preparation. Strand specific cDNA libraries were prepared and sequenced on Illumina NovaSeq6000 platform (2×150bp, paired end) from Eurofins Genomics, Konstanz, Germany.

### Bioinformatic analysis of RNA-seq data

The preprocessing, gene quantification and differential gene expression analysis was performed with the ARMOR workflow^91^. In brief, reads were quality filtered and adapters were removed with Trim Galore! (Krueger, F., TrimGalore. Retrieved February 24, 2010, from https://github.com/FelixKrueger/TrimGalore). For visualization purposes, reads were mapped to the mouse reference genome GRCm38 with STAR version 2.4.2a^87^ and default parameters using Ensembl gene annotations (version 90). BAM files were converted to BigWig files with bedtools^92^. Transcript abundance estimates were computed with Salmon version 0.10.2^93^ and summarized to gene level with the tximeta R package^94^. All downstream analyses were performed in R and the edgeR package^88^ was used for differential gene expression analysis. We filtered the lowly expressed genes and kept all genes with a CPM of at least 10/median_library_size*1e6 in 4 replicates (the size of the smallest group, here the nuclear samples). Additionally, each kept gene is required to have at least 15 counts across all samples. The filtered set of genes was used for the PCA plot and differential gene expression analysis.

### cDNA synthesis and Quantitative Real-Time PCR

Total RNA was reverse transcribed using Superscript III kit (Invitrogen). For qRT-PCR, 2x SYBR master mix (Thermoscientific) were used and the reaction was run in Thermocycler (Applied Biosystems ViiA 7) following the manufacturer’s instructions.

### Primer list

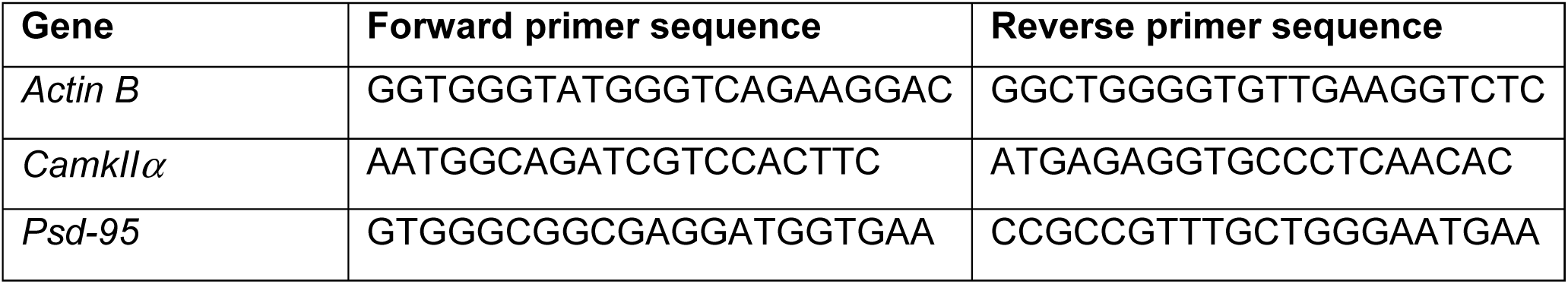

### SDS-PAGE and Western blotting

Protein concentrations were determined using the Pierce BCA Protein Assay (Thermo Fisher Scientific) prior to SDS-PAGE. 20 μg for total protein were used for western blots. The samples were resuspended in 1X SDS loading buffer with 1X final sample reducing reagent and boiled at 95°C, 10 mins. Samples were separated by Bolt 4-12% Bis-Tris pre-cast gels and transferred onto nitrocellulose membranes using iBlot® transfer NC stacks with iBlot Dry Blotting system (Invitrogen). Membranes were blocked with buffer containing 0.05% v/v Tween-20 (Sigma P1379) prepared in PBS (PBST) with 5% w/v non-fat skimmed powdered milk and probed with primary antibodies (list attached) overnight at 4°C in PBST with 1% w/v milk. Following three washes with PBST, membranes were incubated with secondary HRP-conjugated goat anti mouse or rabbit AffiniPure IgG antibodies (1:5000, 1:10000, respectively) (Jackson ImmunoResearch 115-035-146 and 111-035-144, respectively) in PBST with 1% w/v milk, for 1.5 hours at RT. Membranes were washed with PBST, and the bands were visualized using Amersham Imager 600RGB (GE Healthcare Life Sciences 29083467).

### Transmission Electron Microscopy

SNS pellets were prepared from cortical tissue of 1-month-old C57/Bl6 mice as previously mentioned before and submitted to imaging facility at ZMB UZH. Briefly, SNS pellet prepared were re-suspended in 2X fixative (5% Glutaraldehyde in 0.2 M Cacodylate buffer) and fixed at RT for 30 mins. Sample was then washed twice with 0.1 M Cacodylate buffer before embedding into 2% Agar Nobile. Post-fixation was performed with 1% Osmium 1 hour on ice, washed three times with ddH_2_O, dehydrated with 70% ethanol for 20 mins, followed by 80% ethanol for 20 mins, 100% for 30 mins and finally Propylene for 30 mins. Propylene: Epon Araldite at 1:1 were added overnight followed by addition of Epon Araldite for 1 hour at RT. Sample was then embedded via 28 hours incubation at 60°C. The resulting block was then cut into 60 nm ultrathin sections using ultramicrotome. Ribbons of sections were then put onto TEM grid and imaged on TEM – FEI CM100 electron microscope (modify).

### Confocal image acquisition and analysis

Confocal images were acquired on a Leica SP8 Falcon microscope using 63X (NA 1.4) with a zoom power of 3. Images were acquired at a 2048×2048 pixel size, yielding to a 30.05 nm/pixel resolution. To quantify the density of synaptic markers, images were acquired in CA1 region in the apical dendrite area, ∼50 μm from the soma, at the bifurcation of the apical dendrite of pyramidal cells, using the same parameters for both genotypes. Images were acquired from top to bottom with a Z step size of 500 nm. Images were deconvoluted using Huygens Professional software (Scientific Volume Imaging). Images were then analyzed as described previously^84^. Briefly, stacks were analyzed using the built-in particle analysis function in Fiji^95^. The size of the particles was defined according to previously published studies^80,96,97^. To assess the number of clusters, images were thresholded (same threshold per marker and experiment), and a binary mask was generated. A low size threshold of 0.01 μm diameter and high pass threshold of 1 μm diameter was applied. Top and bottom stacks were removed from the analysis to only keep the 40 middle stacks. For the analysis, the number of clusters per 40z stacks was summed and normalized by the volume imaged (75153.8 μm3). The density was normalized by the control group. The densities were compared by t test for 1- and 6-month-old mice. GluN1 synaptic localization was analyzed by counting the number of colocalized GluN1 clusters with Synapsin 1. Colocalization clusters were generated using ImageJ plugin colocalization highlighter. The default parameters were applied to quantify the colocalization. The number of colocalized clusters were quantified using the built-in particle analysis function in Fiji^95^.

### Synaptic density and composition imaging and analysis of primary neuronal culture

Imaging and quantification were performed as previously reported^98^. Briefly, synaptic density and synapse composition was assayed in 22 DIV neuronal cell cultures. Cultures were fixed in cold 4% PFA with 4% sucrose for 20 minutes at RT. Primary antibodies were incubated overnight at 4°C. secondary antibodies were incubated for 3h at RT. Hippocampal primary culture: pyramidal cells were selected based on their morphology and confocal images were acquired on a Leica SP8 Falcon microscope using 63X (NA 1.4) with a zoom power of 3 and analyzed with Fiji software. After deconvolution (huygens professional), images were subsequently thresholded, and subsequent analyses were performed by an investigator blind to cell culture treatment.

### Antibody list

| Antibody | Species, Source | STORM dilution | Confocal dilution | Western blot dilution |
| --- | --- | --- | --- | --- |
| FUS | Rb, A300-293A, Bethyl |  | 1:500 | 1:1000 |
| FUS | Rb, A300-294A, Bethyl |  |  | 1:1000 |
| FUS | Ms, 4H11, Santa Cruz | 1:200 |  |  |
| PSD-95 | Ms, Invitrogen | 1:200 | 1:1000 | 1:1000 |
| P-CAMKIIa | Ms, D21E4, Cell signaling | 1:500 | 1:500 | 1:1000 |
| PNF | Ms, SMI31, Covance |  | 1:1000 |  |
| Spinophilin | Rb, Synaptic Systems | 1:500 |  |  |
| Synapsin 1 | Ms, Synaptic Systems | 1:200 | 1:500 |  |
| GluA1 | Rb, Sigma Aldrich | 1:200 | 1:200 | 1:1000 |
| GluN1 | Ms, Covance |  | 1:500 |  |
| GluN2B | Rb, Sigma Aldrich | 1:500 |  | 1:2000 |
| Bassoon | Gp, Synaptic Systems | 1:500 | 1:500 |  |
| GRP78 BiP (ER) | Rb, Abcam | 1:200 |  |  |
| MAP2 | Ms, Sigma Aldrich |  | 1:1000 |  |
| SYP | Ms, Santa cruz |  |  | 1:200 |
| GABAA/alpha1 | Gp, Synaptic Systems |  | 1:500 |  |
| GABAA/alpha3 | Rb, Synaptic Systems |  | 1:500 |  |
| Gephyrin | Ms, Synaptic Systems |  | 1:500 |  |
| Vgat | Gp, Synaptic Systems |  | 1:500 |  |
| β-Actin | Ms, Sigma | | | 1:5000 |
| SNAP25 | Gp, Synaptic Systems |  | 1:500 | 1:1000 |

## Author Contribution

Conceptualization of the study was carried by S.S., K.M.H., and M.P.. S.S. performed synaptosome isolation, CLIP-seq sample preparation and RNA-seq sample preparation. K.M.H. analyzed the data from CLIP-seq and RNA-seq. S.S., K.M.H., M.D.R. and M.P. developed the strategy to analyze the sequencing data. E.T., M.H.P., M.P.B., J.W., and P.S. provided experimental support for the experiments. L.D. provided the mouse model and input on the study. P.D.R. performed immunostaining and image analyses including confocal, STED and dSTORM. S.S., K.M.H, E.T, P.D.R and M.P wrote and edited the manuscript. M.D.R, P.D.R and M.P. provided supervision. M.P directed the entire study. All authors read, edited, and approved the final manuscript.

## Acknowledgments

We gratefully acknowledge the support of the National Centre for Competence in Research (NCCR) RNA & Disease funded by the Swiss National Science Foundation. SSMK was supported by Swiss Government Excellence Scholarships for Foreign Scholars. The authors would like to thank Prof. Adriano Aguzzi and Dr. Claudia Scheckel for helpful discussions and Dr. Dorothee Dormann for critical comments on the manuscript. We thank Gery Barmettler and Dr. José María Mateos from ZMB UZH for technical help with TEM. We also thank Catharina Aquino and Lucy Poveda from FGCZ for discussions and technical help on CLIP library preparation and sequencing.

## Supplemental Figures titles and legends

**Supplementary Fig. 1 FUS is enriched at the presynaptic compartment**

**(a)** Confocal images showing the distribution of FUS (green) in the molecular layer of the CA1 hippocampal area along with MAP2 (blue) and PNF (magenta). Left panel shows the overview and the right panel, the zoomed in area labelled with the red box on the left panel. **(b)** Similar confocal images showing FUS (green) along with PSD95 (orange) and Synapsin 1 (Syn, blue). (**c**) Schematic of the workflow for distance calculation after STED imaging. (**d**) Schematic of the workflow for distance calculation after STORM imaging. (**e**) Representative images of STORM imaging for FUS-GluN2B-Synapsin1 and FUS-PSD95-Bassoon. (**f**) Violin graph representing the distance distribution between FUS and synaptic markers. (**g**) Binning distribution showing the distance between FUS and the markers (in relative frequency) for PSD95, GluN2b, GluA1, Bassoon, Synapsin and BiP.

**Supplementary Fig. 2 CLIP-seq on cortical synaptoneurosomes identified FUS-associated pre- and postsynaptic RNAs**

(**a**) Western blot of synaptic proteins (GluN2b, SNAP25, GluA1, NRXN1), nuclear protein (Histone H3) in total cortex and synaptoneurosomes (SNS). (**b**) Schematic of CLIP-seq workflow from total homogenate and SNS from mouse cortex. (**c**) Immunoblot showing efficient immunoprecipitation of FUS from total cortex and SNS. (**d**) Flow chart illustrating the reads analyzed to define FUS peaks in total and SNS. (**e**) MA-plot of CLIPper peaks predicted in the total cortex CLIP-seq sample. logCPM is the average log2CPM of each peak in the total cortex and SNS sample and logFC is the log2 fold-change between the number of reads in the total cortex and SNS sample. (**f**) Same MA-plot as (**e**) showing the selected, total cortex specific peaks (p-value cutoff of 3e-03) in red. (**g**) Bar plot of different sets of SNS peaks and their location in genes. The p-value cutoff of each set is on the x-axis and no cutoff refers to the full list of all predicted SNS CLIPper peaks. The selected cutoff is in bold. (**h**) Bar plot of different sets of total cortex peaks and their location in genes. The p-value cutoff of each set is on the x-axis and no cutoff refers to the full list of all predicted SNS CLIPper peaks. The selected cutoff is in bold. (**i**) GO terms enriched among the synapse specific FUS RNA targets.

**Supplementary Fig. 3 Increased synaptic FUS localization in *Fus*^*ΔNLS/+*^ mice affect GABAergic synapses**

(**a**) Western blot of total FUS, full length FUS and actin in synaptoneurosomes isolated from 1-month-old *Fus*^*+/+*^ and *Fus*^*ΔNLS/+*^ mice. (**b**) Quantification of total FUS and full length FUS levels in synaptoneurosomes from *Fus*^*+/+*^ and *Fus*^*ΔNLS/+*^ at 1 month of age. (**c**) Confocal images of the hippocampal CA1 area from 1-month-old mice showing higher level of FUS in the dendritic tree and synaptic compartment in *Fus*^*ΔNLS/+*^ mouse-model. On the top, low magnification pictures show the dendritic area of pyramidal cells stained with FUS (green), MAP2 (dendritic marker, magenta), Synapsin 1 (Syn, Synaptic marker, Cyan) and DAPI (Blue). Red box indicates the area imaged in the high magnification images below. (**d**) Higher magnification equivalent to the area highlighted in red in (**c**). (**e**) Workflow for synaptic marker quantification. Molecular layer of CA1 hippocampal area was imaged by confocal microscopy. Z-stacks were imaged from top (higher Z step with specific signal) to bottom (last step with specific signal) with a Z-step of 0.5 μm. The 40 middle steps were used for quantification. Confocal images were then processed with Huygens professional software for deconvolution. Fiji was used for quantification. Images were first thresholded to only select the specific signal. Images were then binarized and quantification of size and density of synaptic markers was performed using the built-in “Analyze particles”, with size exclusion threshold (as described in the Method section). Data were then compiled in open-office and analyzed using Graphpad Prism software. (**f**) Heatmap summarizing the density of the different synaptic markers quantified in the CA1 hippocampal area from 1-month-old *Fus*^*ΔNLS/+*^ mice. Densities were normalized by the respective control. Mean value of each marker is indicated. Shade of color code for mean variation from 0 (white) to 2 (dark blue). *p<0.05. (**g**) Heatmap summarizing the density of the different synaptic markers quantified in the CA1 hippocampal area from 6-month-old *Fus*^*ΔNLS/+*^ mice. Densities were normalized by the respective control (*Fus*^*+/+*^). Mean value of each marker is indicated. Shade of color code for mean variation from 0 (white) to 2 (dark blue). *p<0.05. (**h**) Heatmap summarizing the cluster area of the different synaptic markers quantified in the CA1 hippocampal area from 1-month-old *Fus*^*+/+*^ and *Fus*^*ΔNLS/+*^ mice. Mean value of each marker is indicated. Shade of color code for mean variation from 0.01 (white) to 1 (dark red). *p<0.05. (**i**) Heatmap summarizing the cluster area of the different synaptic markers quantified in the CA1 hippocampal area from 6-month-old *Fus*^*+/+*^ and *Fus*^*ΔNLS/+*^ mice. Mean value of each marker is indicated. Shade of color code for mean variation from 0.01 (white) to 1 (dark red). *p<0.05 **p<0.01.

**Supplementary Fig. 4 Age-dependent alterations in the synaptic RNA profile of *Fus*^*ΔNLS/+*^ mouse cortex.**

(**a**) Overlap between transcripts expressed in SNS RNA-seq and expressed genes in forebrain synaptic transcriptome reported previously^99^. Expressed genes are all genes with > 10 reads in 2/3 of the replicates (as defined previously^99^). (**b**) Plot of the first and second principal component of all RNA-seq samples and all expressed genes. The genotype is indicated by the symbol and the preparation and age by the color: 1-month-old mice in light and 6-month-old mice in dark colors. (**c**) Plot of the first and third principal component of all RNA-seq samples. (**d**) GO terms enriched among the significantly upregulated genes at 6 months of age in synaptoneurosomes of *Fus*^*ΔNLS/+*^ compared to *Fus*^*+/+*^. (**e**) Gene ontology (GO) terms enriched among the significantly increased RNAs at 6 months of age in synaptoneurosomes of *Fus*^*ΔNLS/+*^ compared to *Fus*^*+/+*^ (**f**) Heatmap from the set of up- and downregulated genes between total cortex samples from *Fus*^*ΔNLS/+*^ and *Fus*^*+/+*^ at 6 months of age. Genes are on the rows and the different total cortex samples on the columns. The color scale indicates the log2FC between the CPM of each sample and mean CPM of the corresponding *Fus*^*+/+*^ samples at each time point [sample logCPM – mean (logCPM of *Fus*^*+/+*^ samples)]. (**g**) Volcano plots showing the log2 fold change of each gene and the corresponding −log10 (FDR) of the differential gene expression analysis comparing total cortex from *Fus*^*ΔNLS/+*^ to *Fus*^*+/+*^ at 1 month (left panel) and 6 months (right panel) of age. The horizontal line marks the significance threshold of 0.05. Significantly downregulated genes are highlighted in green, upregulated genes in purple.

**Supplementary Fig. 5. FUS peak locations on presynaptic and transsynaptic FUS RNA targets altered in *Fus*^*ΔNLS/+*^ mice.**

CLIP-traces showing FUS binding on (**a**) *Syp* (**b**) *Robo2* (**c**) *Sv2a* (**d**) *Syt1* (**e**) *Chl1* (**f**) *App* (**g**) *Aplp2*

**Supplementary Figure 6. FUS peak locations on postsynaptic FUS RNA targets altered in *Fus*^*ΔNLS/+*^ mice.**

CLIP-traces showing FUS binding on (**a**) *Gria2* (**b**) *Gria3* (**c**) *Atp1a1* (**d**) *Atp1a3* (**e**) *Atp1b1* (**f**) *Spock1* (**g**) *Spock2* (**h**) *Clstn1*

**Supplementary Figure 7. FUS binding on *Gabra1* RNA.**

CLIP-traces showing FUS binding to the long 3’UTR containing isoform of *Gabra1*

