## Supplementary Figures for "Synaptic accumulation of FUS triggers age-dependent misregulation of inhibitory synapses in ALS-FUS mice"

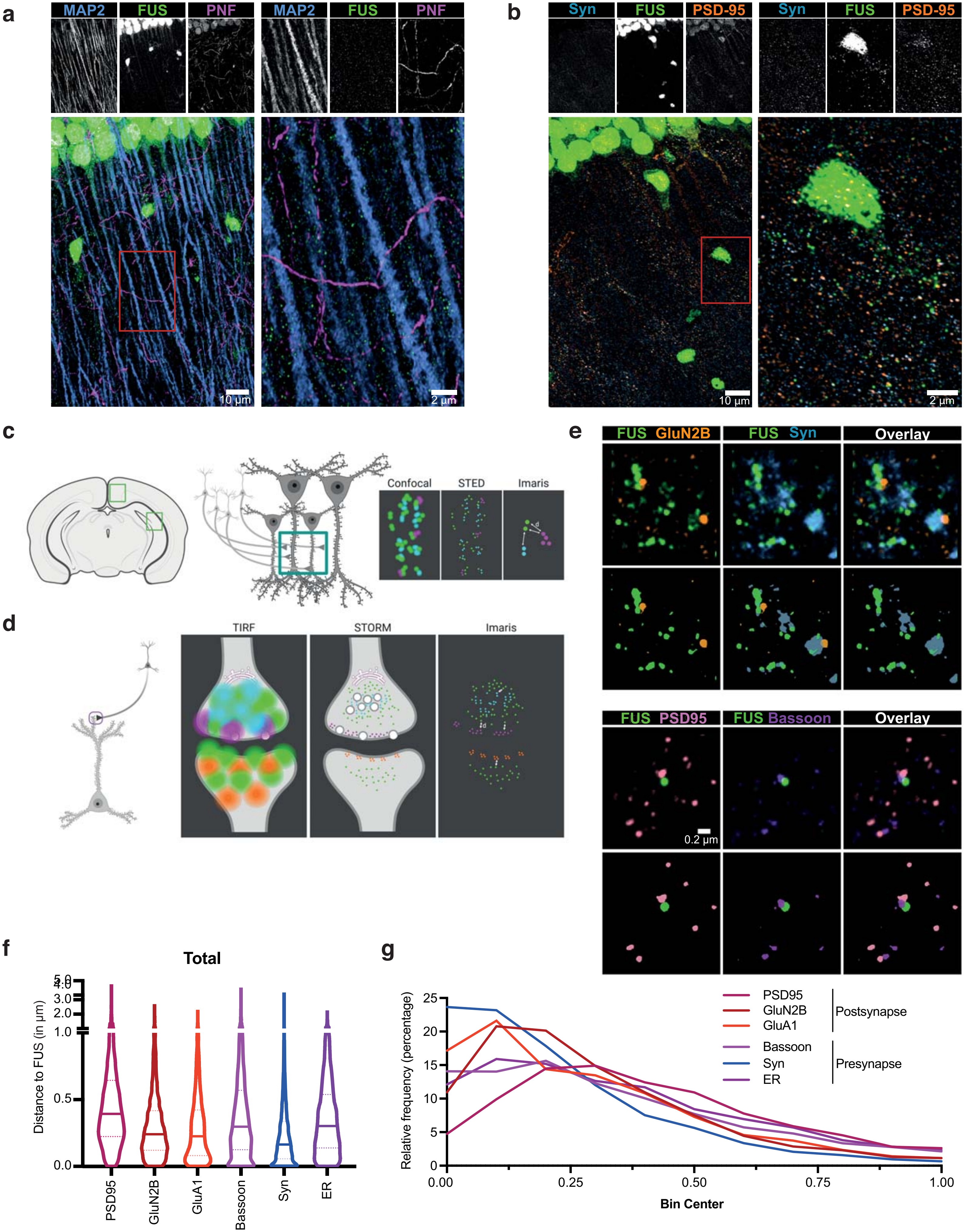


**Supplementary Fig. 1 FUS is enriched at the presynaptic compartment**

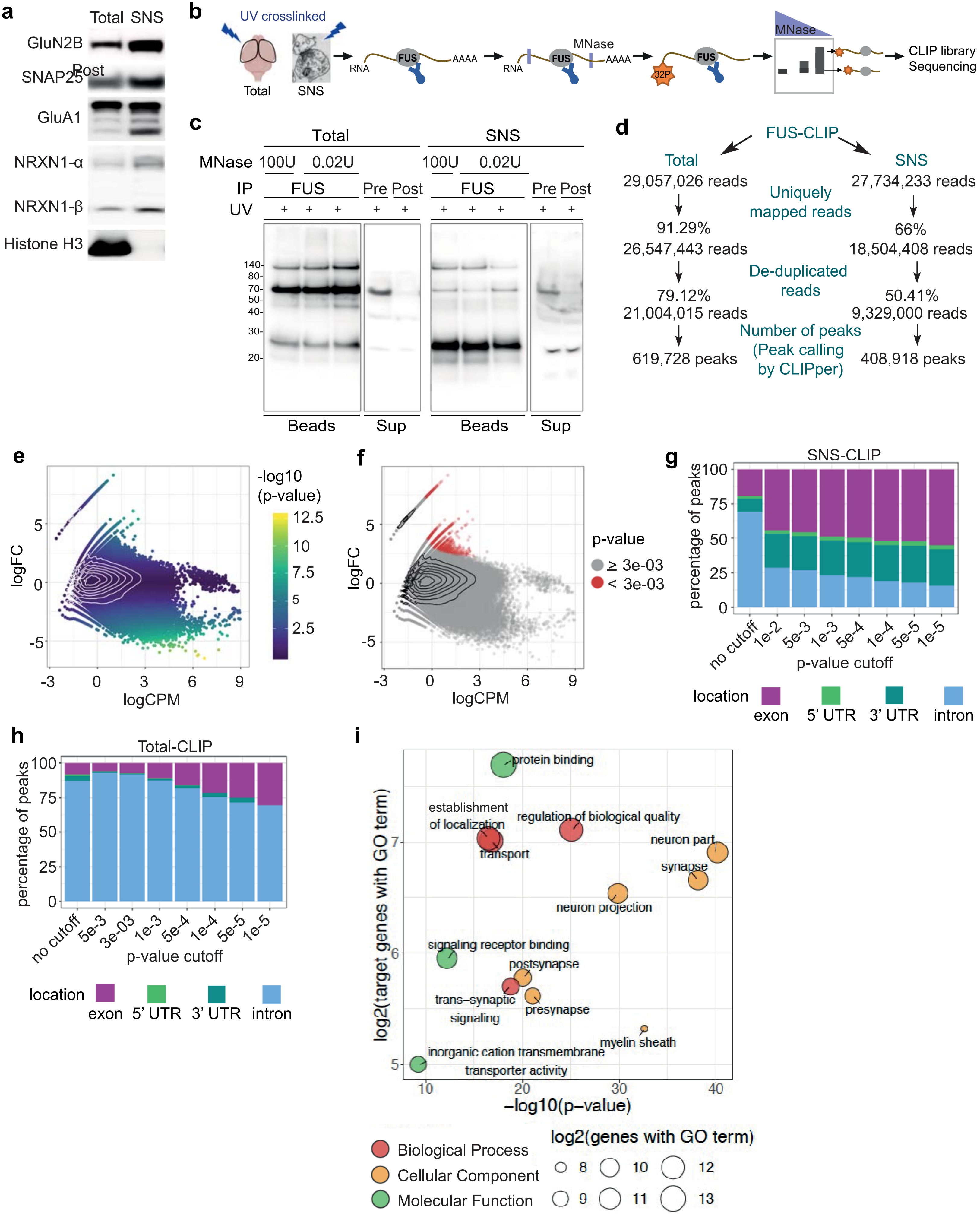


**Supplementary Fig. 2 CLIP-seq on cortical synaptoneurosomes identified FUS-associated pre- and postsynaptic RNAs**

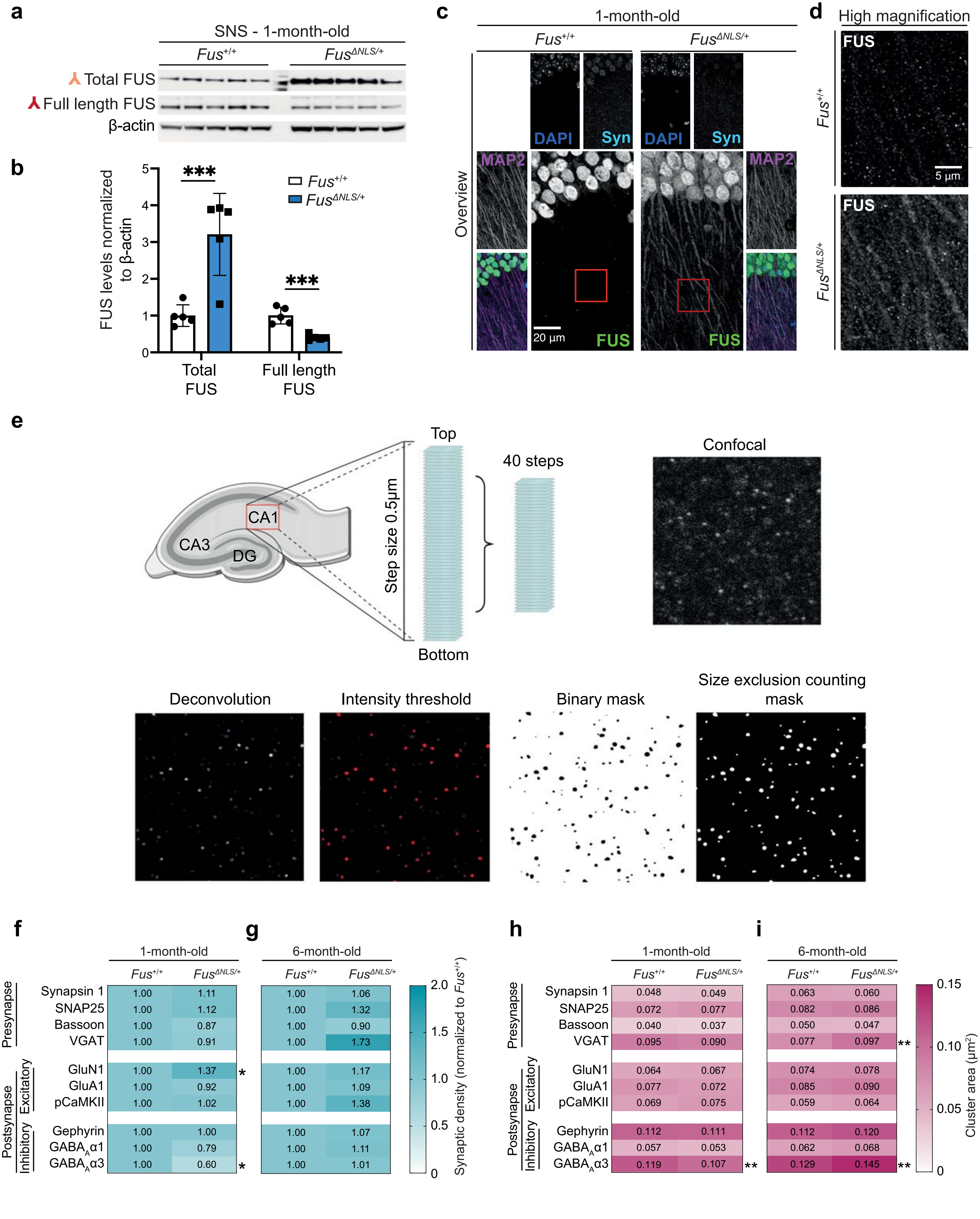


**Supplementary Fig. 3 Age-dependent alterations in the synaptic RNA profile of *Fus*$^{\Delta NLS/+}$ mouse cortex**

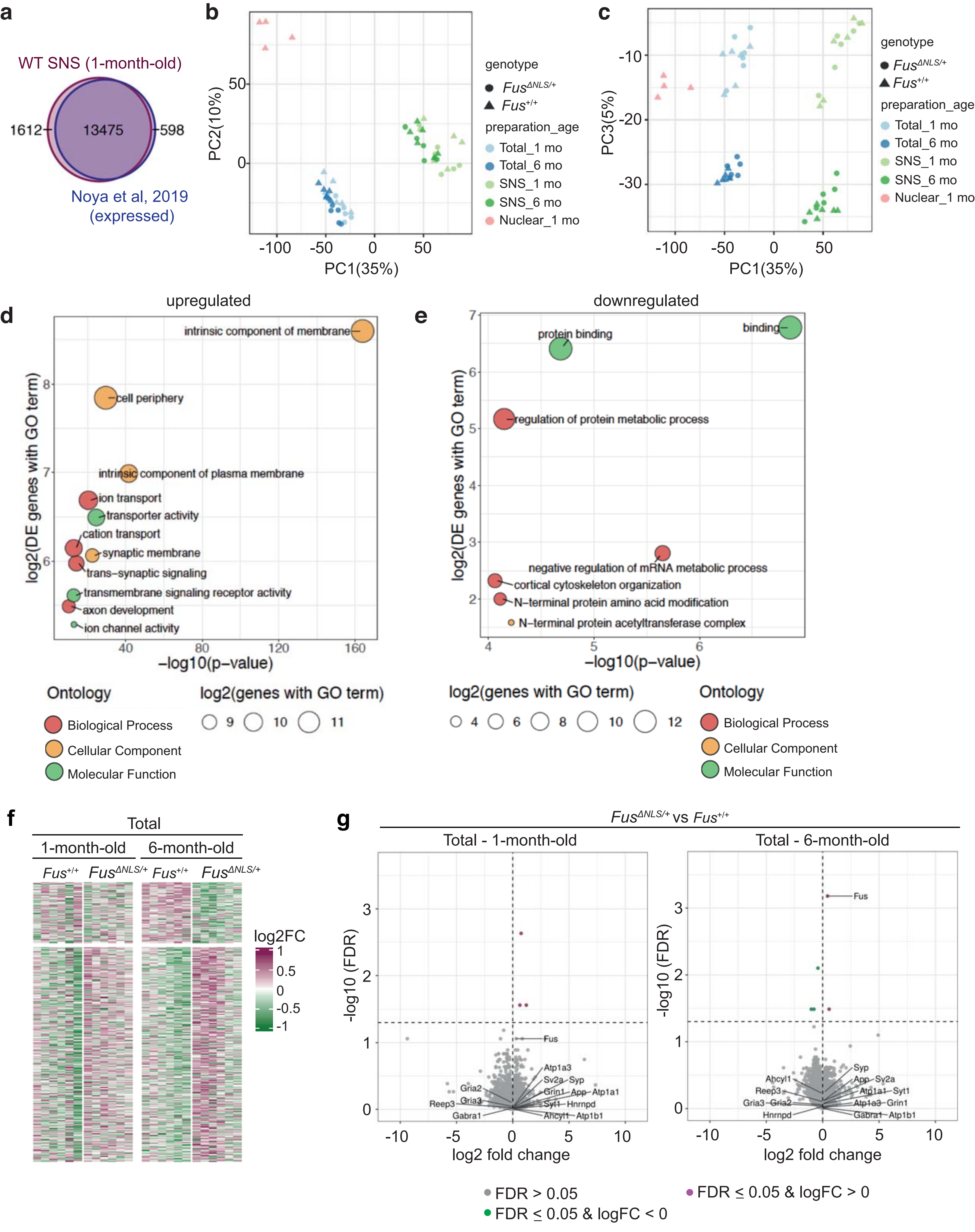


**Supplementary Fig. 4 Age-dependent alterations in the synaptic RNA profile of *Fus*$^{\Delta NLS/+}$ mouse cortex**

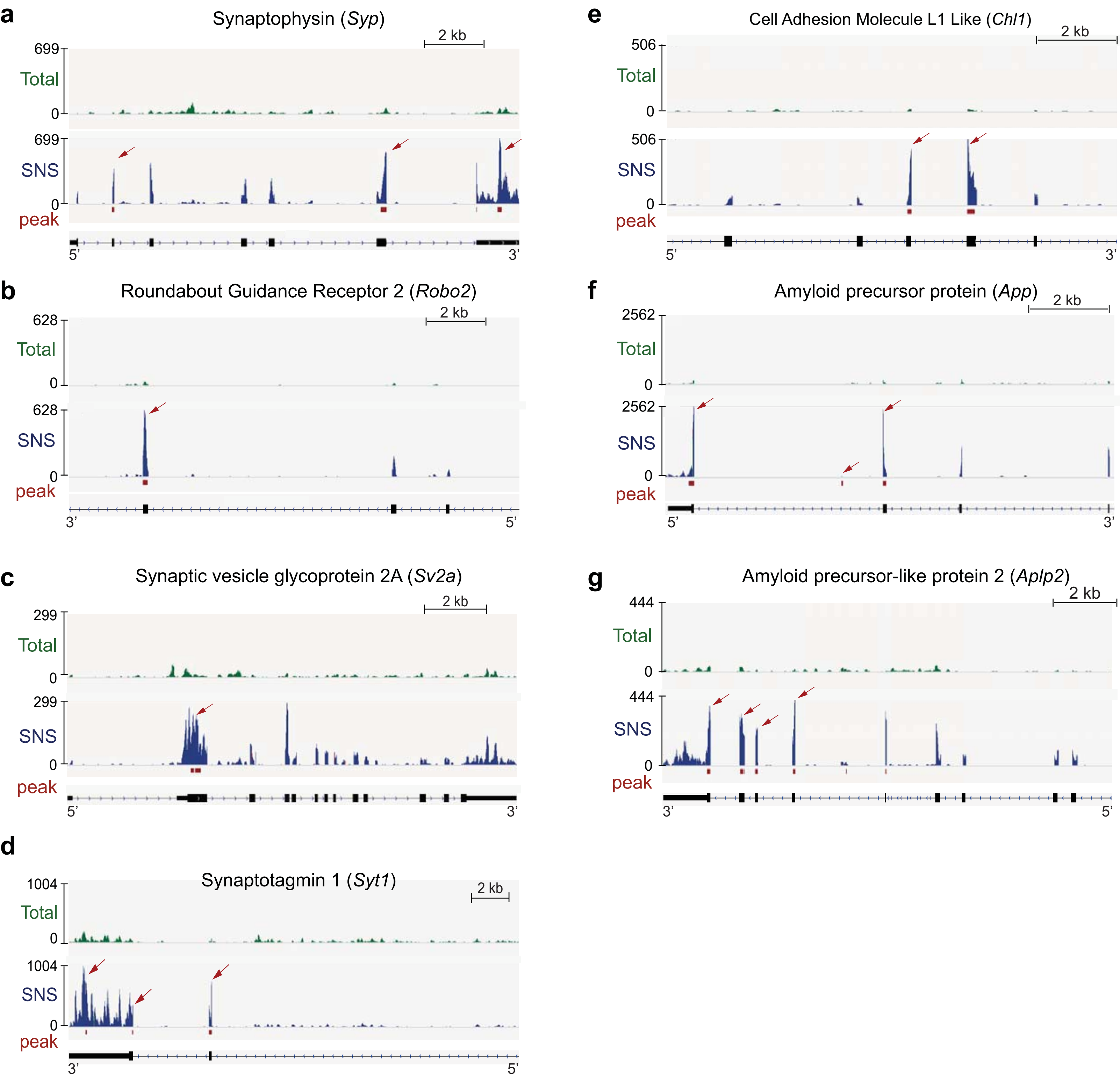


**Supplementary Fig. 5 FUS peak locations on postsynaptic FUS RNA targets altered in *Fus*$^{\Delta NLS/+}$ mice**

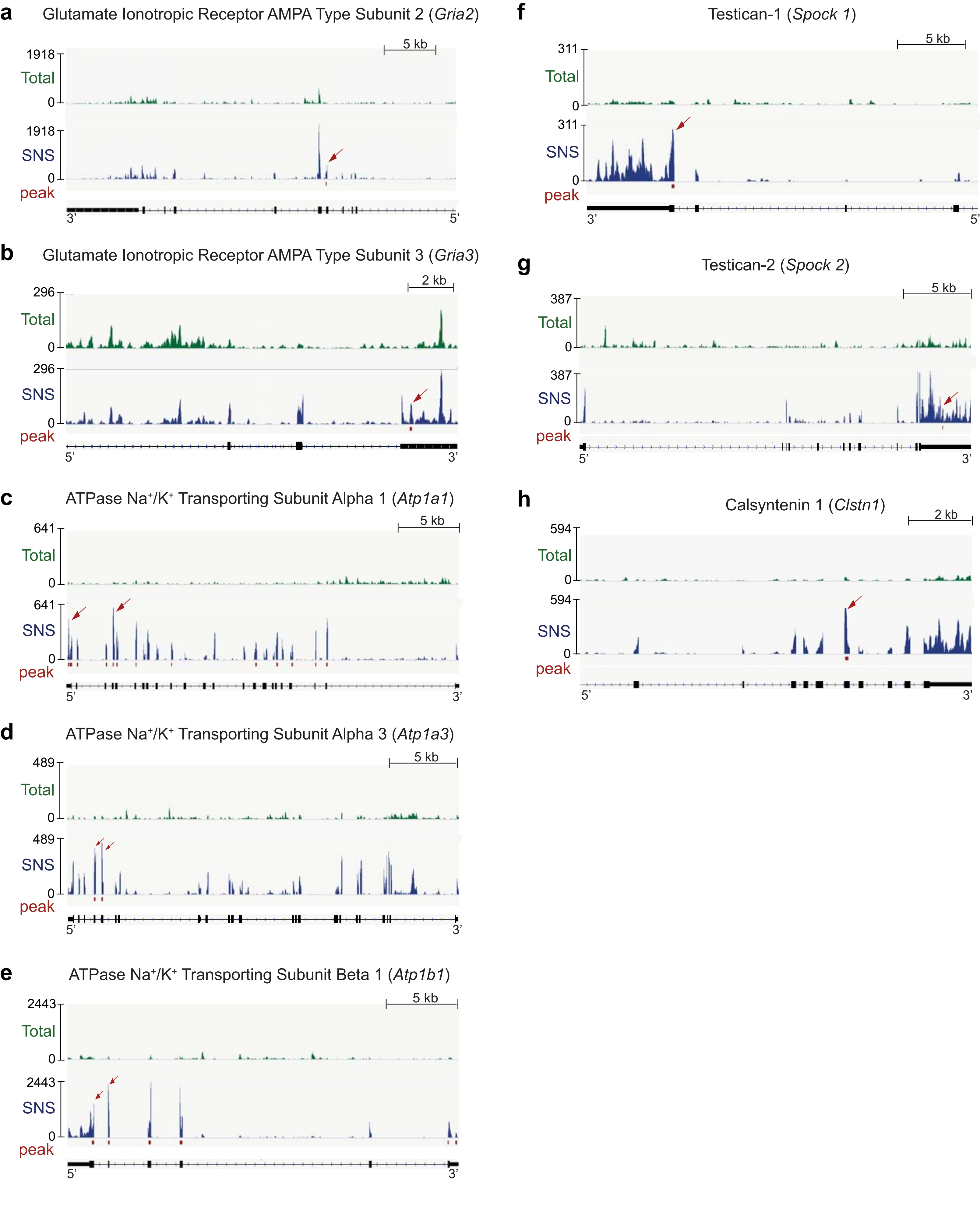


**Supplementary Fig. 6 FUS peak locations on postsynaptic FUS RNA targets altered in *Fus*$^{\Delta NLS/+}$ mice**

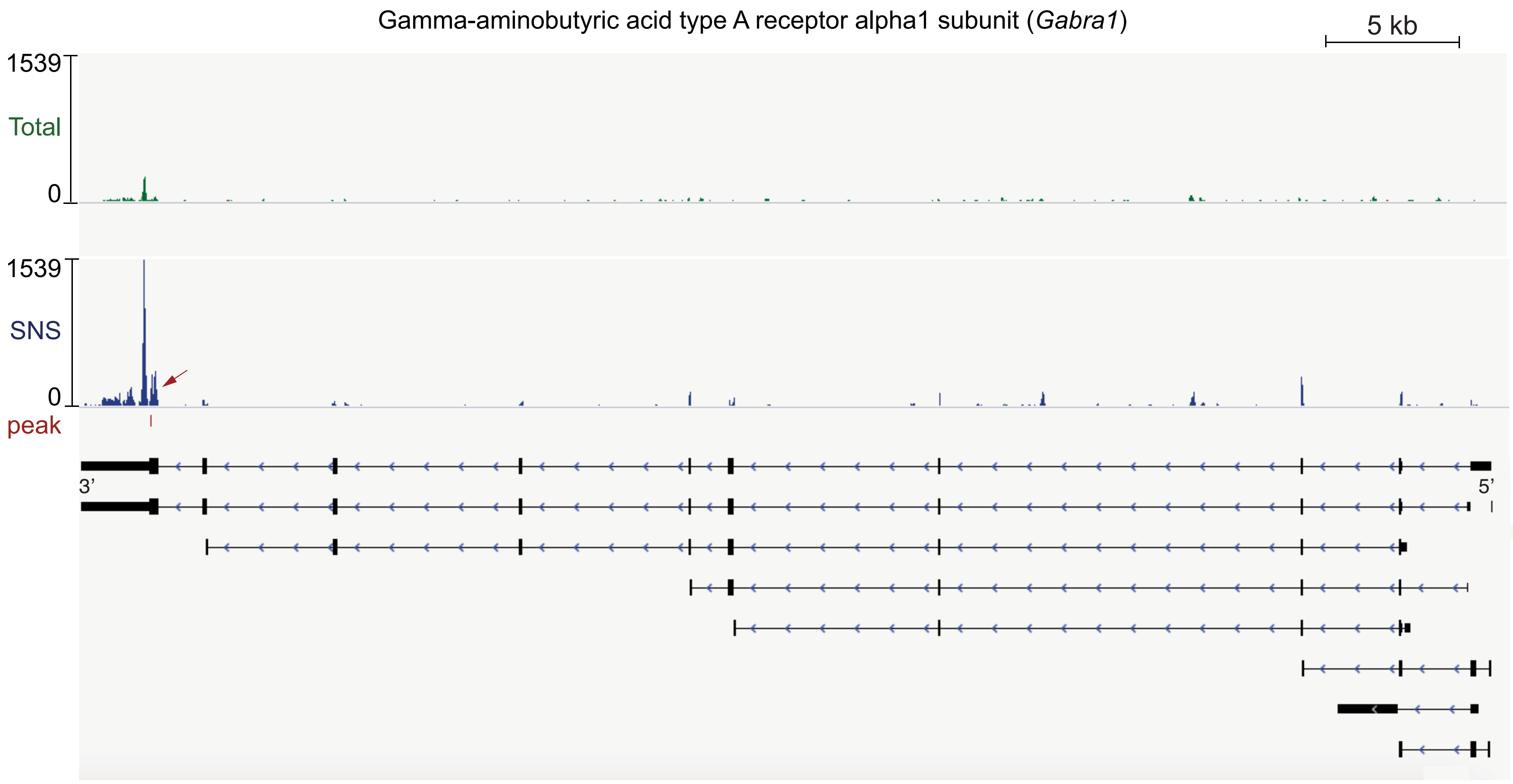


**Supplementary Fig. 7  FUS binding on *Gabra1* RNA**
